# Psychoactive plant- and mushroom-associated alkaloids from two behavior modifying cicada pathogens

**DOI:** 10.1101/375105

**Authors:** Greg R. Boyce, Emile Gluck-Thaler, Jason C. Slot, Jason E. Stajich, William J. Davis, Tim Y. James, John R. Cooley, Daniel G. Panaccione, Jørgen Eilenberg, Henrik H. De Fine Licht, Angie M. Macias, Matthew C. Berger, Kristen L. Wickert, Cameron M. Stauder, Ellie J. Spahr, Matthew D. Maust, Amy M. Metheny, Chris Simon, Gene Kritsky, Kathie T. Hodge, Richard A. Humber, Terry Gullion, Dylan P. G. Short, Teiya Kijimoto, Dan Mozgai, Nidia Arguedas, Matt T. Kasson

## Abstract

Entomopathogenic fungi routinely kill their hosts before releasing infectious spores, but select species keep insects alive while sporulating, which enhances dispersal. Transcriptomics and metabolomics studies of entomopathogens with post-mortem dissemination from their parasitized hosts have unraveled infection processes and host responses, yet mechanisms underlying active spore transmission by Entomophthoralean fungi in living insects remain elusive. Here we report the discovery, through metabolomics, of the plant-associated amphetamine, cathinone, in four *Massospora cicadina*-infected periodical cicada populations, and the mushroom-associated tryptamine, psilocybin, in annual cicadas infected with *Massospora platypediae* or *Massospora levispora*, which appear to represent a single fungal species. The absence of some fungal enzymes necessary for cathinone and psilocybin biosynthesis along with the inability to detect intermediate metabolites or gene orthologs are consistent with possibly novel biosynthesis pathways in *Massospora*. The neurogenic activities of these compounds suggest the extended phenotype of *Massospora* that modifies cicada behavior to maximize dissemination is chemically-induced.

## Introduction

Many animal parasites including viruses, horsehair worms, protists, and fungi modulate the behavior of their hosts, and thereby increase disease transmission (***Berdoy et al., 2000; Gryganskyi et al., 2017; Hoover et al., 2011; Thomas et al., 2002***). Each parasite possesses adaptive traits that maximize dispersal, and often the modification of host behavior is interpreted as a dispersal-enhancing “extended phenotype” of the parasite (***Dawkins, 1982; Hughes et al., 2013***). “Summit disease” (SD) behavior, is an extended phenotype of multiple species of insect-infecting (entomopathogenic) fungi, where parasitized insects ascend and affix to elevated substrates prior to death, which facilitates post-mortem dissemination of spores later emitted from their mummified carcasses (***Elya et al., 2018; Gryganskyi et al., 2017; Hughes et al., 2016; Roy et al., 2006***). A more rarely seen extended phenotype among entomopathogenic fungi is “active host transmission” (AHT) behavior where the fungus maintains or accelerates “normal” host activity during sporulation, enabling rapid and widespread dispersal prior to host death (***Hughes et al., 2016; Roy et al., 2006***).

The Entomophthorales (Zoopagomycota) are among the most important insect- and non-insect arthropod-destroying fungi and include all known species with AHT behavior (***Roy et al., 2006, Spatafora et al., 2017***). *Massospora* is only one of two known genera where AHT is the only reported form of behavior modification across all known species. *Massospora* contains more than a dozen obligate, sexually transmissible pathogenic species that infect at least 21 species of cicadas (Hemiptera) worldwide (***Ciferri et al., 1957; Kobayashi, 1951; Ohbayashi et al., 1999; Soper, 1963; Soper, 1974; Soper, 1981***). Infectious asexual spores (conidia) are actively disseminated from the posterior end of the infected cicada during mating attempts and flights, which are not diminished despite these conspicuous infections (***Cooley et al., 2018; Soper 1963; Soper, 1974; White et al., 1983; Figure 1; Movie 1***). Instead, hypersexual behaviors are observed in *Massospora*-infected cicadas, where male cicadas, in addition to typical mating behavior, also attract copulation attempts from conspecific males (***Cooley et al., 2018; Murphy and Redden, 2003***). *Massospora* effectively hijacks cicadas, turning them into efficient vectors for conidial transmission (***Cooley et al., 2018***).

**Figure 1.**
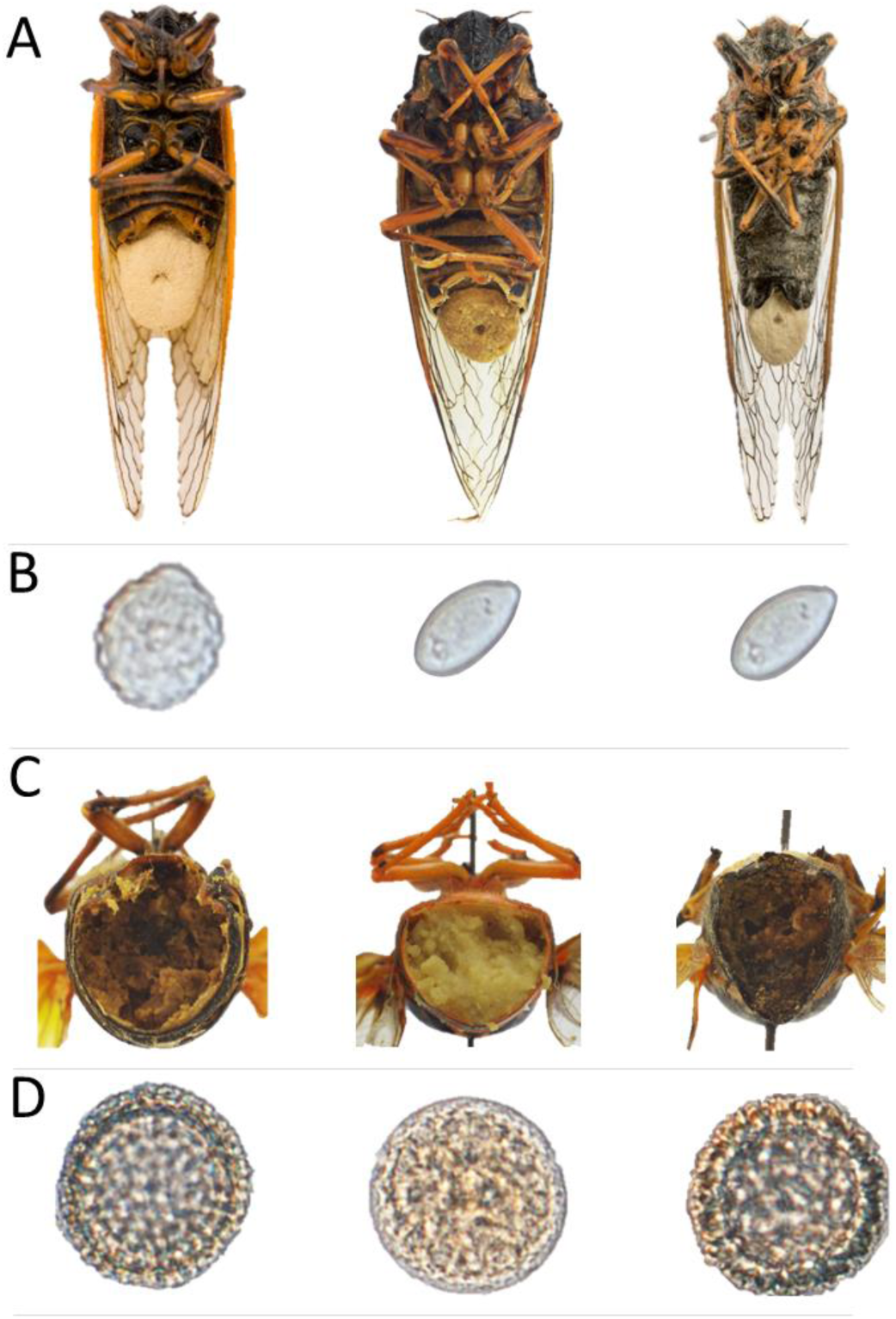
*Massospora*-infected cicadas with associated spore morphology. (A) From left to right: *Mas. cicadina*-infected periodical cicada (*Magicicada septendecim*), *Mas. levispora*-infected Say’s cicada (*Okanagana rimosa*), and *Mas. platypediae* infected wing-banger cicada (*Platypedia putnami*) with a conspicuous conidial “plugs” emerging from the posterior end of the cicada; (B) close-up of conidia for each of three *Massospora* spp.; (C) posterior cross-section showing internal resting spore infection; and (D) close-up of resting spores for each of three *Massospora* spp. Specimens in B-D appear in same order as A.

Because of their ephemeral nature, obligate lifestyle, and large genome size, Entomophthorales with both SD and AHT behaviors have been grossly underrepresented in *in vitro* lab investigations as well as in phylogenetic and phylogenomic studies (***Gryganskyi et al., 2017; Spatafora et al., 2017***). However, recent genomic, transcriptomic, and metabolomic approaches have further helped elucidate important aspects of SD-inducing entomopathogens (***Arnesen et al., 2018; De Fine Licht et al., 2017; Elya et al., 2018; Grell et al., 2011; Malagocka et al., 2015; Wrońska et al., 2018***). Likewise, -omics based investigations of SD-inducing Hypocrealean entomopathogens have helped unravel infection processes and host responses, especially for the “zombie ant fungus” *Ophiocordyceps unilateralis* (***De Bekker et al., 2014; De Bekker et al., 2017; Fredericksen et al. 2017***). This pathogen is thought to modify host behavior via fungus-derived neurogenic metabolites and enterotoxins, thus providing a chemical basis for this behavioral phenotype (***De Bekker et al., 2014; De Bekker et al., 2017***). The mechanisms underlying AHT are completely unexplored, largely due to the inability to axenically culture most Entomophthorales and rear colonies of their insect hosts (such as cicadas) under laboratory conditions. Here, we applied global and targeted liquid chromatography–mass spectrometry (LC-MS)-based metabolomics to freshly collected and archived parasitized cicadas in order to identify candidate small molecules potentially contributing to a hypersexual behavioral phenotype of three *Massospora* species (***Cooley et al., 2018***) or more broadly to AHT (***Batko and Weiser, 1965; Hughes et al., 2016; Hutchison, 1962; Roy et al., 2006***).

## Results

### Morphology of *Massospora* specimens

Morphological studies were conducted to permit species-level identification of collected specimens based on previously published studies (***Soper, 1963; Soper, 1974***). Conidia (n = 37) and resting spore (n = 13) measurements were acquired from freshly collected *Mas. cicadina-* infected periodical cicadas (*Magicicada* spp.) and *Mas. platypediae*-infected wing-banger annual cicadas (*Platypedia putnami*), as well as from archived *Mas. levispora-*infected *Okanagana rimosa and Mas. cicadina-*infected periodical cicadas (***Figure 1, Table 1***). Morphology of conidia were studied in 15, 14, and 8 specimens of *Mas. cicadina* (*Mc*), *Mas. platypediae* (*Mp*), and *Mas. levispora* (*Ml*), respectively, while morphology of resting spores was examined in 11, 1 and 1 specimens (***Table 2, Figure 1–figure supplement 1 & 2***). The conidial specimens of *Mas. cicadina*-infected periodical cicadas spanned eight Broods (populations of periodical cicadas that emerge in the same year and that tend to be geographically contiguous) and six of the seven *Magicicada* species collected over 38 years and distributed across the eastern U.S., while *Mas. platypediae*- and *Mas. levispora*-infected cicadas were from a single annual cicada populations collected in New Mexico (2017) and Michigan (1998), USA, respectively (***Table 2, Figure 1– figure supplement 1 & 2***).

**Table 1.** Sample size for morphological, molecular, metabolomics, and genomic studies of *Massospora*-infected cicadas and their asymtomatic counterparts. Sample origin refers to the authors/lab who collected specimens. Numbers before the slash denote conidial isolates whereas those after denote resting spore isolates. Bracketed numbers denote isolates for which sequences were deposited in NCBI Genbank.

| Cicada host/<br>population | *Massospora* sp. | Year<br>collected | Sample<br>origin | Morphology | Phylogenetics | Global<br>Metabolomics | Targeted<br>Metabolomics | Genomics |
| --- | --- | --- | --- | --- | --- | --- | --- | --- |
| *Magicicada septendecula* Brood I (WV, USA) | *Mas.cicadina* | 1978 | JRC/CS | 1/0 | 1/0 | – | – | – |
| *Magicicada* spp. Brood II (NY & CT, USA) | *Mas.cicadina* | 1979 | JRC/CS, RAH | 3/0 | 1/0 | – | – | – |
| *Magicicada septendecula* Brood IV or XIX (MO, USA) | *Mas.cicadina* | 1998 | JRC/CS | 1/0 |  |  |  |  |
| *Magicicada* spp. Brood V (OH & WV, USA) | *Mas.cicadina* | 2016 | MTK, NA, WJD | 5/0 | 3/0 [1] | 5/0 | 20/0 | 1/0 |
| *Magicicada septendecim* Brood VI (NC, USA) | *Mas.cicadina* | 2017 | TG | 0/8 | 0/3 | 0/4 | 0/4 | – |
| *Magicicada septendecim* Brood VIII (PA, USA) | *Mas.cicadina* | 2002 | JRC/CS | 1/0 | – | – | 0/5 | – |
| *Magicicada cassini* Brood IX (NC, USA) | *Mas.cicadina* | 2003 | JRC/CS |  |  |  | 0/1 |  |
| *Magicicada cassini* Brood X (OH, USA) | *Mas.cicadina* | 2004 | GK | 1/0 | 1/0 | – | – | – |
| *Magicicada septendecula* Brood XIV (OH, USA) | *Mas.cicadina* | 2008 | GK | 1/0 | 1/0 | – | – | – |
| *Magicicada septendecula* Brood XIX (IL, USA) | *Mas. cicadina* | 1998 | JRC/CS | 0/3 | 1/0 |  |  |  |
| *Magicicada tredecim* Brood XXII (MS, USA) | *Mas. cicadina* | 2001 | JRC/CS | – | – | – | 3/0 |  |
| *Magicicada* spp. Brood XXIII (IL & IN, USA) | *Mas. cicadina* | 1989, 1998, 2002, 2015 | JRC/CS, GK | 2/0 | 1/0 [1] | – | 0/4 | – |
| *Okanagana rimosa* (MI) | *Mas. levispora* | 1998 | JRC/CS | 8/1 | 3/0 [1] | – | 7/1 | – |
| *Platypedia putnami* (NM) | *Mas. platypediae* | 2017 | MTK | 14/1 | 3/0 [1] | 5/0 | 11/0 | 1/0 |
| *Magicicada* spp. Brood V | None | 2016 | MTK | – | – | 2 | 2 | – |
| *Okanagana rimosa* (MI) | None | 1998 | JRC/CS | – | – | – | 1 | – |
| *Platypedia putnami* (NM) | None | 2017 | MTK | – | – | 1 | 1 | – |
| *Magicicada* sp. Brood VII | None | 2018 | MTK | – | – | – | 10 | – |
| *Platypedia putnami* (CA) | None | 2018 | MTK | – | – | – | 10 | – |
| **Total** |  |  |  | **50** | **18** | **17** | **80** | **2** |

**Table 2.** Mean (A) conidia and (B) resting spore dimensions for three *Massospora* species sampled from infected cicadas. Twenty-five conidia or resting spores were measured for each specimen except for *Mas. levispora* (MI) and *Mas.* aff. *levispora* (NM) resting spores, in which 50 spores were measured.

| *Massospora* sp. | Reference | N | Conidia | N | Resting spores |
| --- | --- | --- | --- | --- | --- |
| *Mas. cicadina* | Soper 1976 | | 10 – 17 $\mu\text{m}$ x 14 – 20 $\mu\text{m}$ (avg. 15.5 x 18.5 $\mu\text{m}$) | | 33.5 $\mu\text{m}$ x 47.5 $\mu\text{m}$ (avg. 40.5 $\mu\text{m}$) |
| | This study | 15 | 12.9 – 18.1 $\mu\text{m}$ x 13.2 – 18.9 $\mu\text{m}$ (15.4 x 16.0 $\mu\text{m}$) | 11 | 41.7 $\mu\text{m}$ x 43.2 $\mu\text{m}$ (avg. 42.2 $\mu\text{m}$) |
| *Mas. levispora* (MI) | Soper 1963 | | 6.0 – 11.0 $\mu\text{m}$ x 9.5 – 23.0 $\mu\text{m}$ (8.0 x 15.0 $\mu\text{m}$) | | (avg. 34.0 $\mu\text{m}$) |
| | This study | 8 | 7.5 – 10.2 $\mu\text{m}$ x 11.5 – 18.3 $\mu\text{m}$ (8.8 x 14.3 $\mu\text{m}$) | 1 | 38.8 $\mu\text{m}$ x 40.7 $\mu\text{m}$ (avg. 39.8 $\mu\text{m}$) |
| *Mas. aff. levispora* (NM) | Soper 1976 | | 5.2 – 7.7 $\mu\text{m}$ x 6.5 – 12.9 $\mu\text{m}$ (avg. 6.0 x 9.9 $\mu\text{m}$) | | not previously reported |
| | This study | 14 | 6.5 – 9.6 $\mu\text{m}$ x 9.9 – 17.5 $\mu\text{m}$ (8 x 13.2 $\mu\text{m}$) | 1 | 40.5 $\mu\text{m}$ x 41.4 $\mu\text{m}$ (avg. 40.9 $\mu\text{m}$) |

Mean conidia (*Mc* = 15.4 µm x 16.0 µm and *Ml* = 8.8 µm x 14.3 µm) and resting spore (*Mc* = 42.4 µm and *Ml* = 39.8 µm) dimensions for *Mas. cicadina* and *Mas. levispora* overlapped with previously reported measurements (***Table 2, Figure 1–figure supplement 1 & 2***). In contrast, spore dimensions (8 x 13.2 µm) in the majority of *Mas. platypediae* specimens fell completely outside the reported range for this species (***Table 2, Figure 1–figure supplement 1***). Interestingly, all the *Mas. platypediae* measurements fell within the reported range for *Mas. levispora*, including a single resting spore specimen, which had not been previously observed for *Mas. platypediae* (***Table 2, Figure 1–figure supplement 2***).

### Phylogenetic relationships among *Massospora* morphotypes

To infer evolutionary relationships among collected specimens, phylogenetic analyses were performed. Prior to this study, *Massospora* was represented in NCBI Genbank by a total of four DNA sequences from a single isolate (ARSEF 374) of *Mas. cicadina* (NCBI Genbank accessions EF392377, EF392548, EF392492, and DQ177436). A total of 18 isolates representing *Mas. cicadina* (n = 12), *Mas. levispora* (n = 3), and *Mas. platypediae* (n = 3) were used for single gene phylogenetic studies, four of which were included as representatives isolates in the combined phylogeny (***Table 1***). Reference LSU, SSU, and EFL DNA sequences for *Mas. cicadina, Mas. platypediae* and *Mas. levispora* (MI) were deposited in NCBI Genbank (MH483009 - MH483011, MH483015-MH483023 and MH547099).

Phylogenetic analysis of the partitioned individual alignments (data not shown) and combined dataset (LSU+SSU) using maximum likelihood (ML) resolved all sampled *Massospora* in a strongly supported monophyletic group sister to a clade containing *Entomophthora* and *Arthrophaga* (***Figure 2***). EFL was excluded from the concatenated dataset on account of its heterogeneous distribution (some taxa have EF1 and others have EFL; see ***James et al., 2006***) across the Entomophthorales taxa included in the phylogenetics analysis. However representative sequences were deposited into Genbank for all three *Massospora* species (see above). *Massospora cicadina* isolates formed a single lineage that encompassed all sampled cicada broods and *Magicicada* species (data not shown). The previously deposited *Mas. cicadina* ARSEF 374 was one exception as it aligned with *Entomophthora*, indicating a wrongly assigned species to previously deposited DNA sequences in Genbank. To confirm this finding, we successfully resampled and amplified the SSU region (deposited as Genbank accession MH483023) from the physical specimen associated with ARSEF 374, which was 100% identical to the other *Mas. cicadina* sequences from this study. *Mas. levispora* and *Mas. platypediae* did not form distinct clades; instead, they appeared as a single lineage sister to *Mas. cicadina* (***Figure 2***). Coupled with the morphological data, we propose two possible scenarios: 1) *Mas. levispora* and *Mas. platypediae* comprise a single annual-cicada infecting *Massospora* species, which occupies a broader geographic and host range than previously reported, or 2) *Mas. levispora* and *Mas. platypediae* are two related species capable of infecting wing-banger cicadas. Because more sampling is needed to resolve these phylogenetic findings, subsequent mentions of *Mas. platypediae* from cicada host *Platypedia putnami* from New Mexico (NM), USA will be referred to hereafter as *Mas.* aff. *levispora* (NM), while *Mas. levispora* sensu stricto from its cicada host *Okanagana rimosa* in Michigan (MI), USA will retain the designation of *Mas. levispora* (MI).

**Figure 2.**
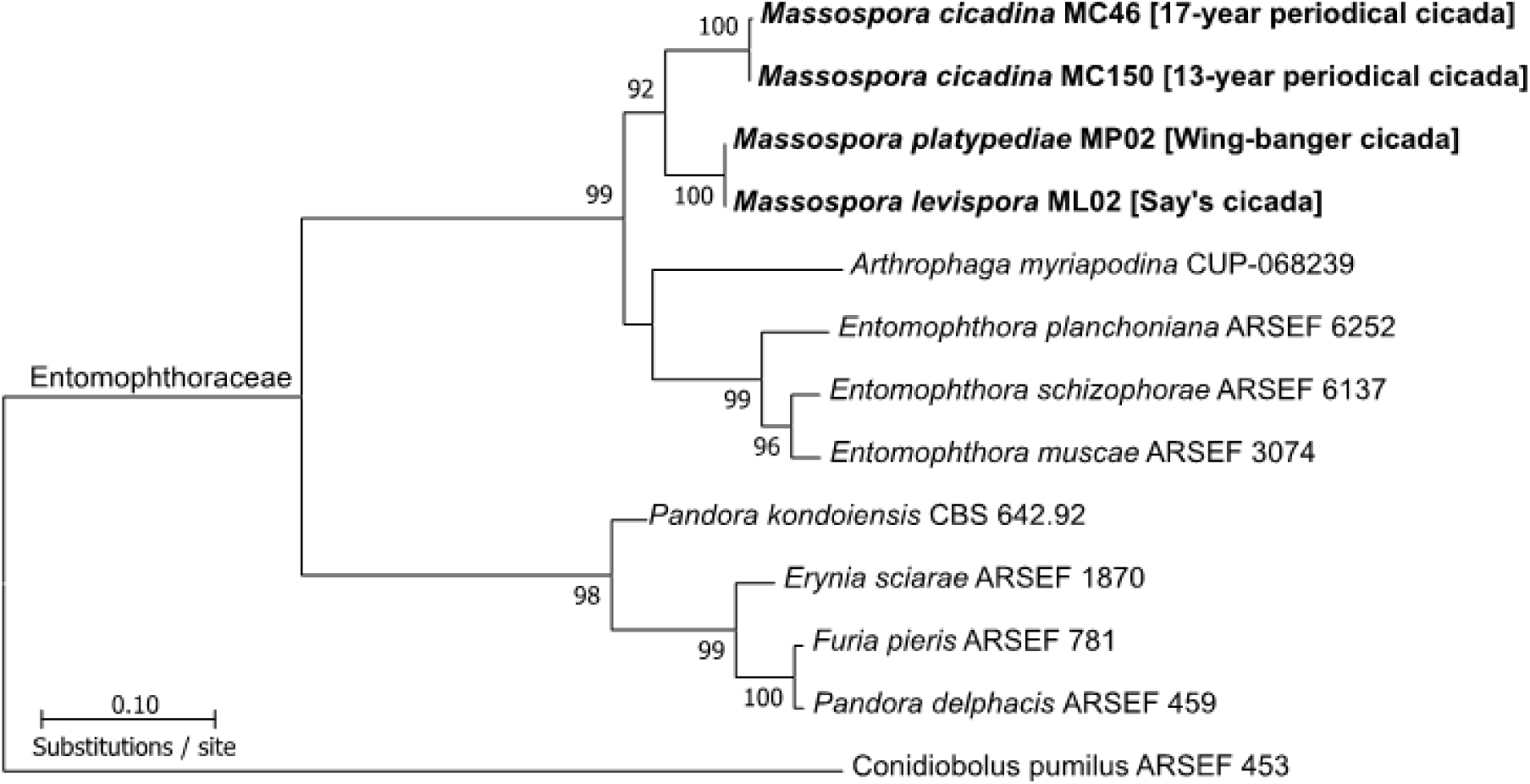
Concatenated LSU+SSU maximum likelihood (ML) tree consisting of *Massospora* species and related species in the Entomophthorales. Given that *Mas. levispora* and *Mas. platypediae* were not genealogically exclusive, *Mas. platypediae* will be hereafter referred to as *Mas.* aff. *levispora* (NM). Bootstrap support is indicated near each node and only values greater than 70% are shown.

### Discovery of monoamine alkaloids using global metabolomics

LC-MS-based global metabolomics using high resolution, accurate mass Quadrupole Time of Flight Liquid Chromatography Mass Spectrometry (QToF LC/MS) was performed on conidial plugs (=exposed mass of conidia and host tissue) excised from *Mas. cicadina*-infected periodical cicadas (*Magicicada* spp.; ***Figure 1A***) and *Mas.* aff. *levispora* (NM)-infected wing-banger cicadas (*Platypedia putnami*; ***Figure 1A, Table 1***). A second global comparison was also performed on conidial and resting spore plugs from *Mas. cicadina*-infected periodical cicadas (***Table 1***). Posterior abdominal segments from healthy cicadas of both species were also sampled as controls (***Table 1***). Each of the three sample types produced unique metabolome profiles, while those of conidial plugs from the two *Massospora* species were more similar compared to the cicada control. In total, 1,176 (678 positive and 489 negative ion mode) small molecule spectra (free amino acids, phospholipids, ceramides, alkaloids, etc.) were differentially detected between the two *Massospora* spp. including two notable monoamine alkaloids.

The amphetamine cathinone (*m/z* 150.0912), previously reported from the plant *Catha edulis* (***Groves et al., 2015***) was detected only in the *Mas. cicadina* samples and not in the *Mas.* aff. *levispora* (NM) conidial plugs nor the healthy cicada controls (***Figure 3***). Cathinone was also detected in resting spore specimens in a second global metabolomics study between *Mas. cicadina* resting spores, an independently collected population sampled from North Carolina (NC), USA in 2017, and the initial conidial plugs from West Virginia (WV), USA in 2016. However, the conidial plugs contained 4.61 ± 0.28 higher fold change compared to resting spores (***Figure 3–figure supplement 1)***. Fragmentation patterns for cathinone in *Mas. cicadina* conidial and resting spore plugs matched the experimentally observed fragmentation patterns of a commercially available DEA-exempt analytical standard used in this study; three fragments with *m/z* less than 10 ppm mass error of those obtained from the analytical standard were recovered (***Figure 4A-C***). Several cathinone pathway intermediates, including 1-phenylpropane-1,2-dione, the metabolite immediately preceding cathinone, were putatively identified from the global metabolomics output but fragmentation patterns did not fall within the accepted mass error limits when compared to experimentally observed fragmentation patterns of a commercially available analytical standard. Other pathway intermediates, initially identified from the global metabolomics output, including benzaldehyde, were not explored further due to their involvement in several divergent biosynthetic pathways (***Lomascolo et al., 1999***).

**Figure 3.**
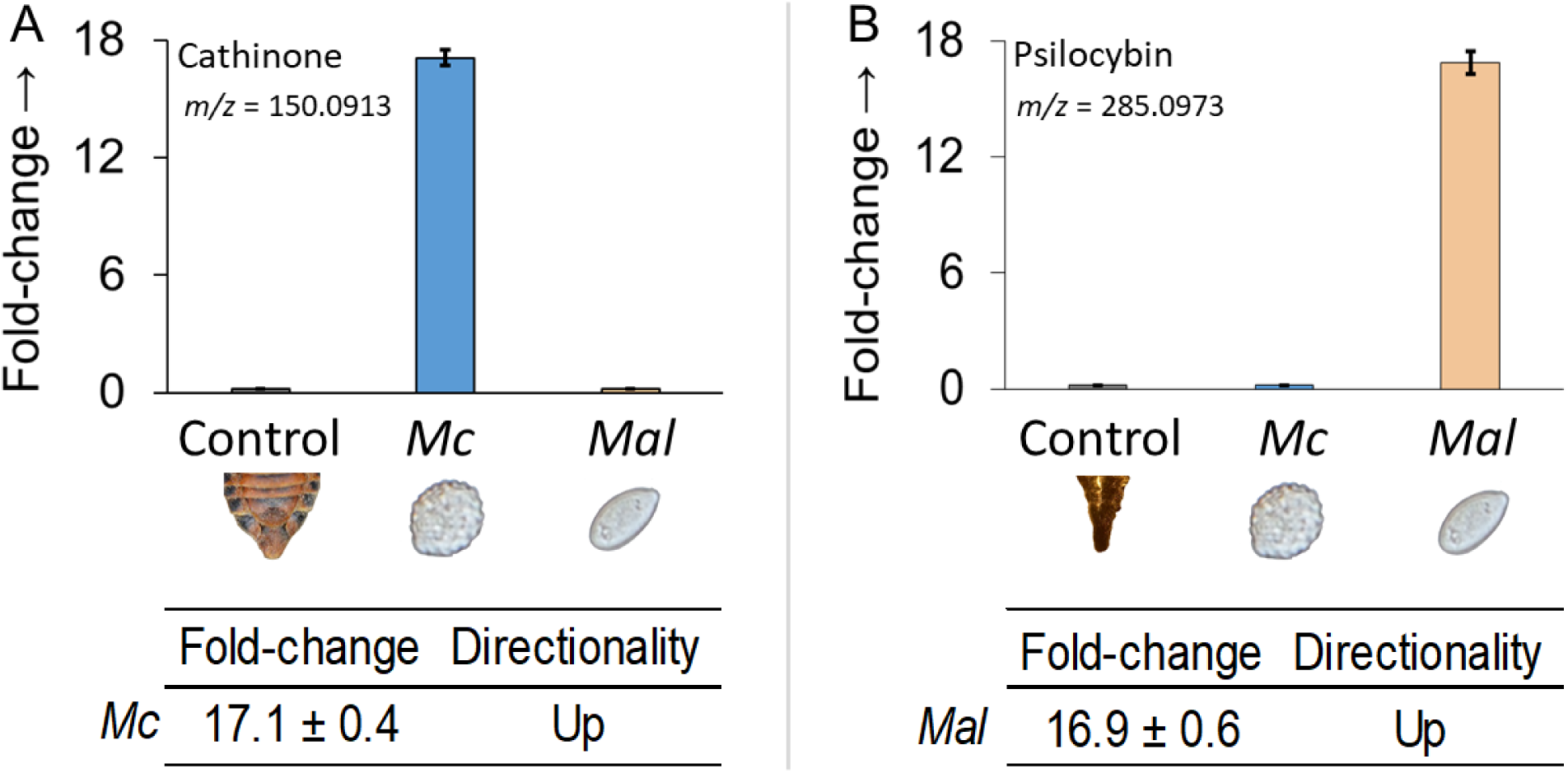
Global metabolomics fold-change comparisons of (A) cathinone and (B) psilocybin among healthy cicada controls (posterior sections), *Mas. cicadina* (*Mc*), and *Mas.* aff. *levispora* (NM) (*Mal*) conidial plugs. Fold change ± standard error based on comparisons with control. Value is average of five biological replicates.

**Figure 4.**
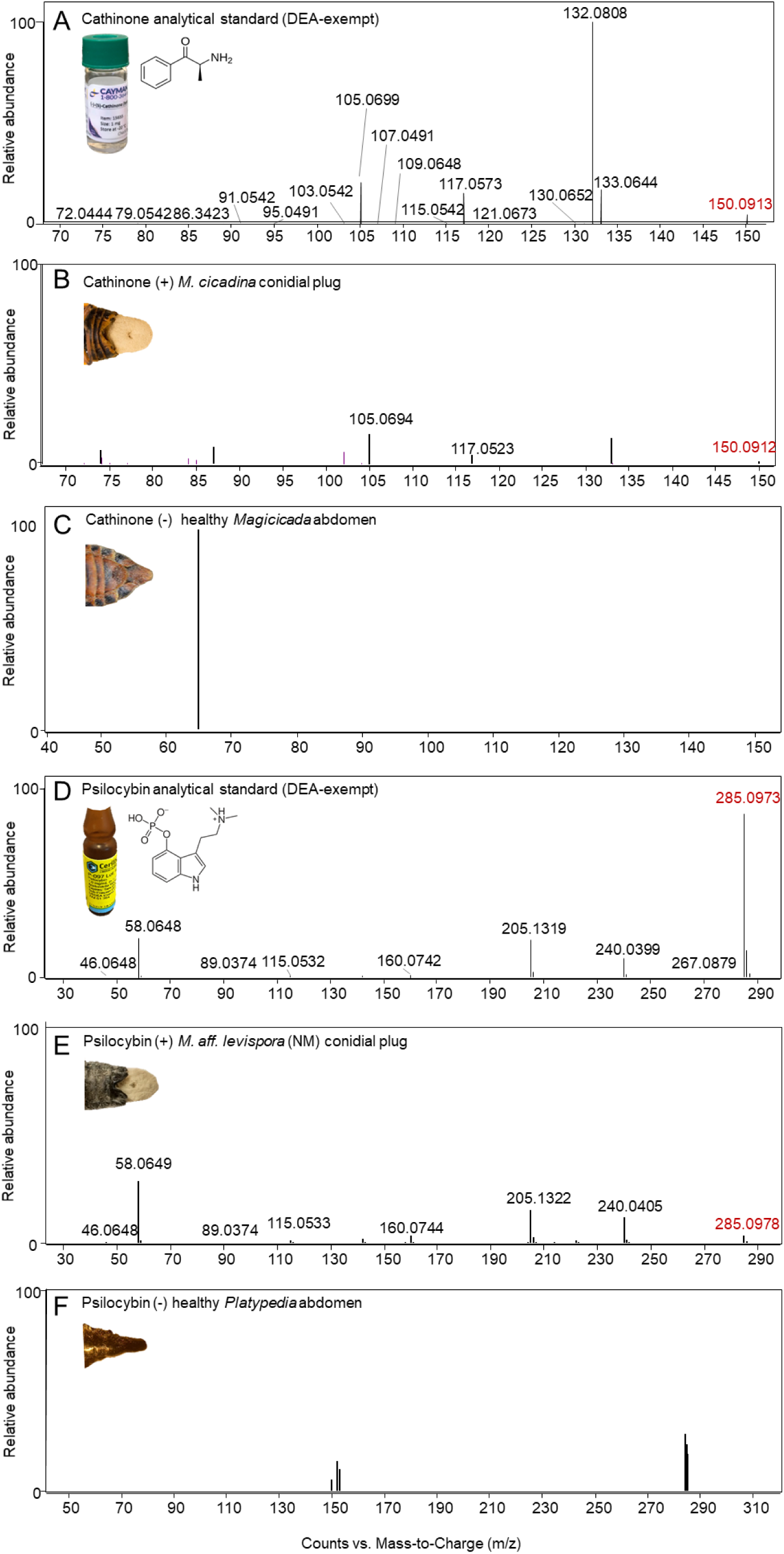
Representative LC-MS/MS spectra for A) cathinone DEA-exempt analytical standard B) cathinone-positive *Mas. cicadina* conidial plugs from infected *Magicicada* spp., B) cathinone-negative *Magicicada septendecim* abdominal sample, D) psilocybin DEA-exempt analytical standard, E) psilocybin-positive *Mas.* aff. *levispora* (NM) conidial plugs from infected *Platypedia putnami*, F) psilocybin-negative *Platypedia putnami* abdominal sample. *M/z* in red denote precursor fragment while *m/z* in black denote observed fragments. All labeled *m/z* values from *Massospora* plugs (B and E) are within 10 ppm mass error compared to analytical standard.

Cathinone CoA-dependent pathway intermediates were also monitored using a targeted method with a triple quadrupole mass spectrometer (QQQ-MS) system. Over 10 CoA species were monitored including all five hypothesized cathinone CoA-dependent pathway intermediates, Trans-cinnamoyl-CoA, 3-Hydroxy-3-phenylpropionyl-CoA, 3-oxy-3-phenylpropionyl-CoA, benzoyl-CoA (***Groves et al. 2015***). The most abundant CoAs detected were Acetyl-CoA and Malonyl-CoA, yet none of the cathinone CoA-dependent pathway intermediates monitored were detected (***data not shown***).

Psilocybin (*m/z* 285.0973), a psychoactive tryptamine produced by about 200 Agaricalean mushroom species (***Kosentka et al., 2013***), was among the most abundant metabolites detected in *Mas.* aff. *levispora* (NM) conidial plugs but was not detected in *Mas. cicadina* or the cicada controls (***Figure 3***). Fragmentation patterns for psilocybin in *Mas.* aff. *levispora* (NM) plugs matched the experimentally observed fragmentation patterns of a commercially available DEA-exempt analytical standard used in this study; eight fragments with *m/z* less than 10 ppm mass error of those obtained from the analytical standard were recovered (***Figure 4D-F***). One intermediate in the psilocybin biosynthetic pathway in *Psilocybe cubensis* mushrooms (***Fricke et al., 2017***), 4-hydroxytryptamine, was identified from the global metabolomics output for both *Mas. cicadina* and *Mas*. aff. *levispora* (NM) but fragmentation patterns could not be compared to an analytical standard since 4-HT is not commercially available for purchase. Analytical stands for other pathway intermediates either don’t exist or lack DEA-exempt formulations.

### Quantification of cathinone, psilocybin, and psilocin using targeted metabolomics

Cathinone, psilocybin, and psilocin were monitored individually using QQQ-MS (***Figure 5A***). We performed targeted LC-MS/MS absolute quantification of cathinone, psilocybin, and psilocybin’s biologically active bioconversion product psilocin, on individual conidial and resting spore plug extracts from all three cicada-fungal pairs (***Table 1***) including *Mas. levispora* (MI)-infected Say’s cicadas (*Okanagana rimosa*), which were excluded from the global metabolomics on account of their previous long-term storage in ethanol. Given its taxonomic overlap with *Mas.* aff. *levispora* (NM), we hypothesized that it might also contain psilocybin. The posterior abdomen of cicada controls used for global metabolomics as well as independently collected healthy *Magicicada* (n = 10) and *Platypedia* (n = 10) cicadas were assessed for all three compounds. All samples were quantified against standard curves generated from DEA-exempt analytical standards. Cathinone was quantified in three collections of *Mas. cicadina*: one from Brood V (2016) in WV, one from brood VI in NC, USA in 2017, and a third from archived collection spanning four broods (VIII, IX, XXII, and XXIII) collected from 2001-2003 (***Cooley et al., 2018***).

**Figure 5.**
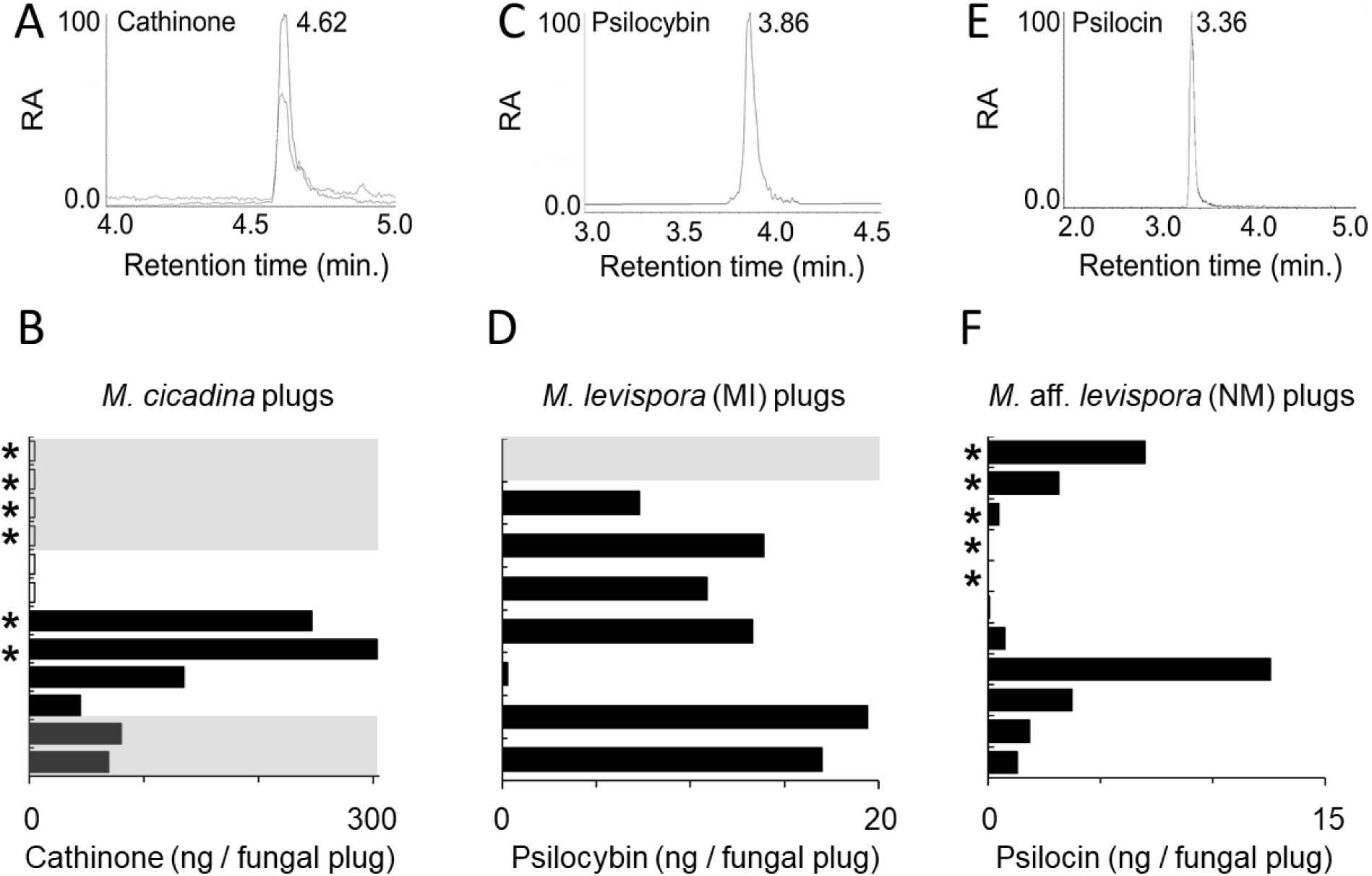
Absolute quantification of (A-B) cathinone, (C-D) psilocybin, and (E-F) psilocin from Massospora fungal plugs using Q-TOF LC/MS. A, C, and E show extracted ion chromatograms with retention times for (A) cathinone with an *m/z* of 150.0913 from *Mas. cicadina* plugs (C) psilocybin with an *m/z* of 285.0973 from *Mas. levispora* (MI) plugs, and (E) psilocin with an *m/z* of 205.1319 from *Mas.* aff. *levispora* (NM) plugs. For cathinone, two different transitions were monitored concurrently. RA denotes relative abundance. B, D, and F show the concentration of each alkaloid for across the same sample sets determined from peak area measured against standard curves generated using DEA-exempt analytical standards. Each sample was injected a single time per run. Each of the three compounds were monitored individually in independent runs. Rows showing short unfilled (white) bars denote the presence of the compound from the sample but concentrations below the limit of quantification. Gray-background denotes samples with resting spore fungal plugs while white background denotes conidial fungal plugs. Asterisk on the far left denote samples used in both global and targeted metabolomics runs. Order of *Massospora* samples from top to bottom as are they appear in ***Figure 5–figure supplement 1***.

Cathinone was present in 4 of 4 Brood VI *Mas. cicadina* resting spore plugs and 6 of 24 Brood V *Mas. cicadina* conidial plugs and was quantifiable in concentrations from 44.3 ? 303.0 ng / infected cicada in four of six Brood V fungal plugs (***Figure 5–figure supplement 1***). From the archived collection, cathinone was present and quantifiable in concentrations of 68.9 and 80.1 ng / infected cicada from 2 of 13 resting spore plugs spanning two separate broods (***Figure 5–figure supplement 1***). A Brood V *Mas. cicadina* conidial plug extraction that had 4-months earlier yielded a total of 246 ng of cathinone and served as a positive control in the absolute quantification of archived samples was found to contain 116 ng, a notable reduction from the previous time point (***Figure 5–figure supplement 1***). This is not unexpected given the labile nature of amphetamines (***Kalix, 1988***).

Psilocybin was detected in 11 of 11 assayed *Mas.* aff. *levispora* (NM) plugs and 7 of 8 assayed *Mas. levispora* (MI) plugs while psilocin was detected in 9 of 11 plugs and 0 of 8 plugs, respectively (***Figure 5***). Of these, psilocybin was quantifiable only in *Mas. levispora* (MI), with concentrations from 7.3 to 19.4 ng / infected cicada across 6 fungal plugs (***Figure 5–figure supplement 1***). Psilocin was detected only in *Mas.* aff. *levispora* (NM) in concentrations from 0.1 to 12.6 µg / infected cicada across 9 fungal plugs (***Figure 5–figure supplement 1***).

To further confirm the fungal origin of monoamine alkaloids from *Massospora*-infected cicadas, we screened independent healthy populations of the two cicada hosts included in the global metabolomics study, *Platypedia putnami* from CA and Brood VII *Magicicada septendecim* from NY, for the presence/absence of cathinone, psilocybin and psilocin. These samples were screened using a high-resolution, accurate mass Orbitrap LC-MS, which allowed for targeted fragmentation of individual alkaloids. Sampling included five outwardly asymptomatic males and females for each cicada species. Similar to data from cicada controls in the global metabolomics study, results of these studies failed to detect any of the three monitored compounds from macerated posterior abdominal sections including their bacteriomes, which in some cicada species have been shown to contain cryptic fungal partners in the secondary-metabolite-rich Hypocreales (***Matsuura et al., 2018; Figure 5–figure supplement 1***).

### Searches for genetic mechanisms of psilocybin and cathinone biosynthesis in *Massospora*

Having confidently detected and quantified cathinone, psilocybin, and psilocin from independently collected *Massospora* populations, we next searched for candidate genes underlying cathinone and psilocybin biosynthesis in assembled *Mas. cicadina* and *Mas.* aff. *levispora* (NM) metagenomes (***Figure 6; Figure 7*).**

**Figure 6.**
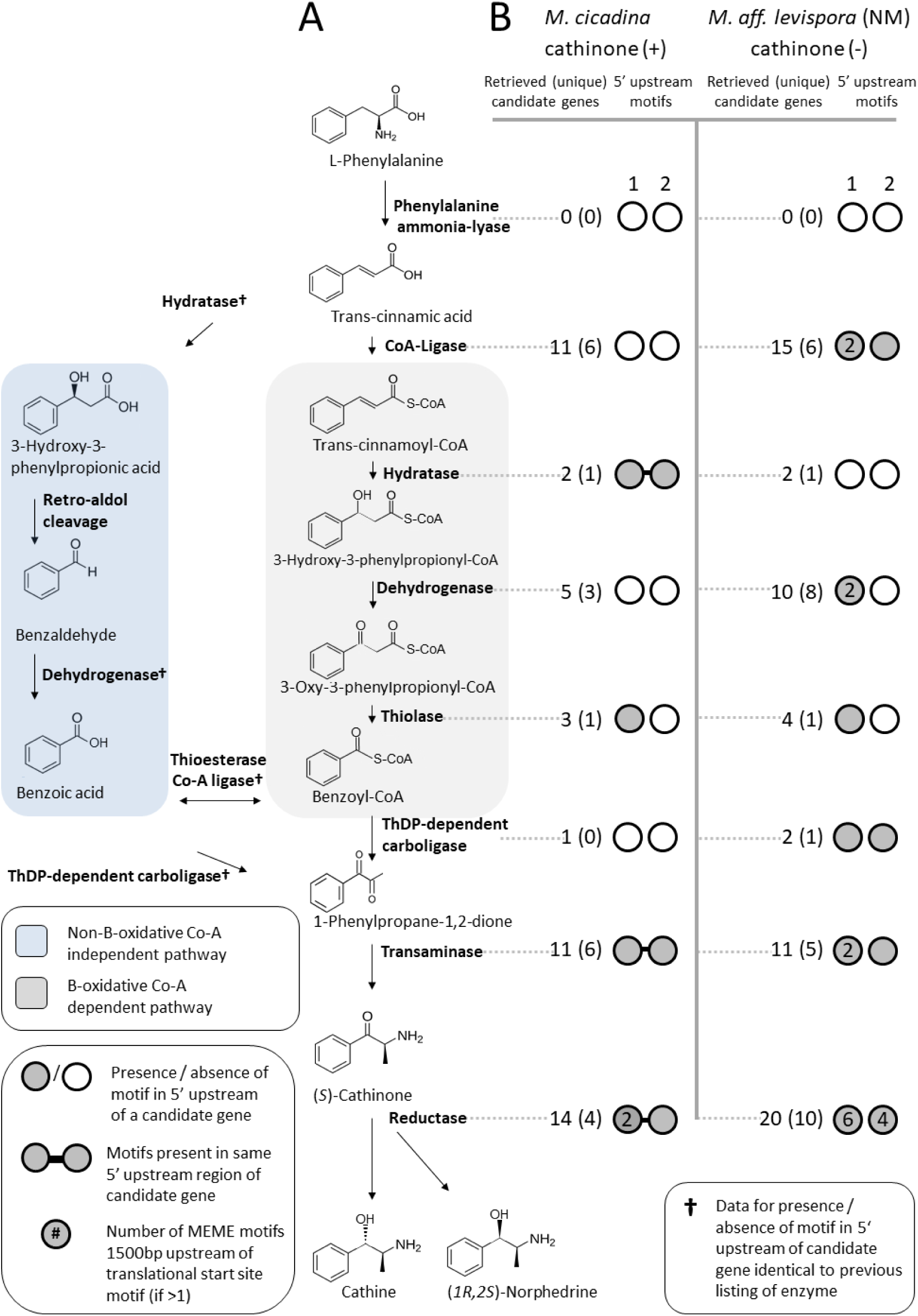
A) Predicted cathinone biosynthesis pathways from *Catha edulis (****Groves et al. 2016***), B) Putative regulatory motifs found upstream of candidate cathinone biosynthesis genes from *Mas. cicadina* and *Mas.* aff. *levispora* (NM) assemblies. Sequences were retrieved using profile HMMs of enzymes predicted for each pathway step. Candidate gene co-regulation was predicted by shared nucleotide motifs within 1500bp upstream of predicted translational start sites. Only motifs associated with transaminase genes (predicted to catalyze the production of cathinone) are shown.

**Figure 7.**
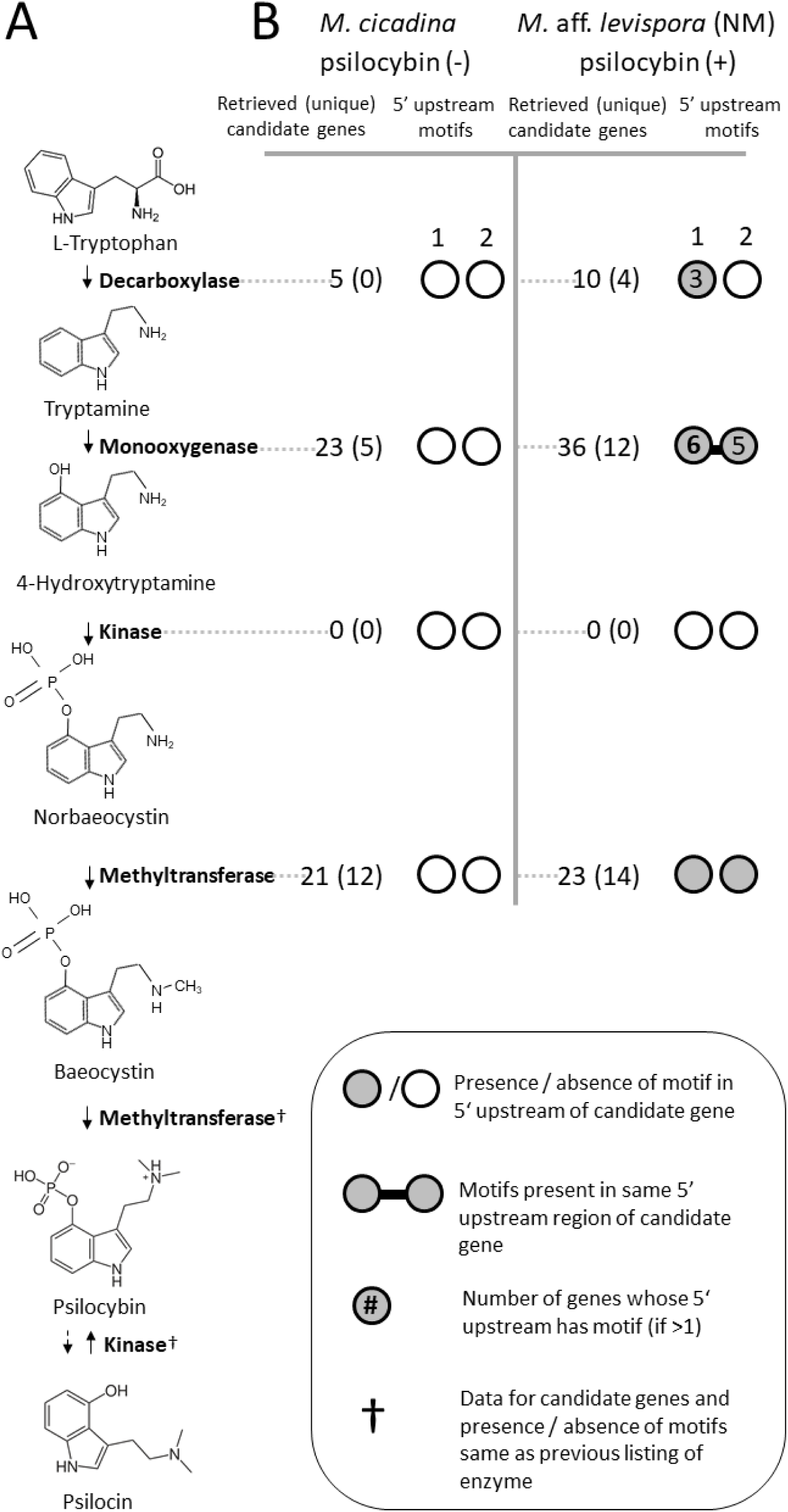
A) Psilocybin biosynthesis pathway from Basidiomycete fungi, B) Putative regulatory motifs found upstream of candidate psilocybin biosynthesis genes from both *Mas. cicadina* and *Mas*. aff. *levispora* (NM) assemblies. Sequences were retrieved using profile HMMs of enzymes predicted for each pathway step. Candidate gene co-regulation was predicted by shared nucleotide motifs within 1500bp upstream of predicted translational start sites. Only motifs associated with methyltransferase genes (which catalyzes the formation of psilocybin) are shown.

We first used tBLASTn searches of each *Massospora* metagenome assembly using four characterized psilocybin biosynthesis proteins from *Psilocybe cubensis* (***Fricke et al., 2017***) and all 42 candidate cathinone biosynthesis proteins identified from RNA sequence data for *Catha edulis* (***Groves et al., 2015***). After failing to retrieve *Massospora* sequences orthologous to genes from the characterized *Psilocybe* psilocybin or predicted *Catha* cathinone pathways in either of our two *Massospora* metagenomes, we developed eight primer sets with various levels of degeneracy in attempt to amplify *Psilocybe* and *Psilocybe*-like psilocybin biosynthesis genes from multiple fungal plugs of *Mas.* aff. *levispora* (NM) in addition to NM-02 from which the metagenomic data was generated (***Table S1***). Fungal plugs from *Mas. levispora* (MI) were also screened for psilocybin biosynthesis genes using the same primer sets. No DNA sequence data were available in GenBank for cathinone biosynthesis (only RNA sequences were deposited in Genbank Short-Read Archive) so primers were not developed. All primer sets failed to amplify Psi-specific PCR targets in any of the assayed *Massospora* DNA templates.

We next searched for all sequences containing protein domains found among the enzymes of each biosynthetic pathway (***Fricke et al., 2017; Groves et al., 2015; Table S2***). In each metagenome, we identified multiple genes representative of all classes of enzymes in each pathway with two exceptions (***Figure 6–figure supplement 1***; ***Table S3-S4***). Although at least one candidate homolog could be identified for 7 of the 8 genes predicted to play a role in cathinone biosynthesis in both the *Mas. cicadina* (cathinone +) and *Mas.* aff. *levispora* (NM) (cathinone -) metagenomes, we failed to detect genes encoding phenylalanine ammonia-lyase (*PAL*) in either metagenome (***Figure 6–figure supplement 2)***. Similarly, in both the *Mas.* aff. *levispora* (NM) (psilocybin +) and *Mas. cicadina* (psilocybin -) metagenomes, we identified at least one candidate homolog for 3 of the 4 genes participating in psilocybin biosynthesis, the exception being 4-hydroxytryptamine kinase (*PsiK*) in either metagenome (***Figure 7–figure supplement 1***). The failure to detect *PAL* and *PsiK* homologs could be attributed to the fragmented and incomplete nature of the *Mas. cicadina* and *Mas.* aff. *levispora* (NM) metagenomes (which had N50 values of 3457bp and 3707bp, respectively), evolution of an alternate enzyme mechanism in *Massospora*, or the execution of these steps either directly or indirectly by the host. Further investigations with higher quality genome assemblies of fungi and hosts are needed to test these hypotheses.

To narrow down our list of candidate genes that may underlie the biosynthesis of these natural products, we employed a two-pronged approach to gather additional lines of evidence that would suggest participation in a common secondary metabolite pathway (***Slot, 2017; Wolf et al., 2015***). First, we looked for evidence of coordinated regulation from shared nucleotide sequence motifs in the upstream non-coding region of each candidate sequence (***Figure 6–figure supplement 1***). We also looked for evidence of gene clustering among homologs of the candidate sequences across a phylogenetically diverse database of 375 fungal genomes, since clustering could not be assessed in the fragmented *Massospora* metagenomes (***Figure 6–figure supplement 1; Table S5***). The two candidate regulatory motifs recovered upstream of candidate cathinone biosynthesis genes in the *Mas. cicadina* assembly (cathinone +) were found upstream of the same predicted hydratase, transaminase and reductase genes (***Figure 6–figure supplement 3, Table S6***). Only two of the seven candidate cathinone regulatory motifs in the *Mas.* aff. *levispora* (NM) assembly (cathinone -) were found upstream of transaminase genes, while the rest were not (***Figure 6–figure supplement 3, Table S6***). Two of five motifs found upstream of candidate psilocybin biosynthesis genes in the *Mas.* aff. *levispora* (NM) assembly (psilocybin +) were present upstream of methyltransferase genes, while the single motif found in the *Mas. cicadina* assembly (psilocybin -) was not (***Figure 7–figure supplement 2, Table S6***).

Searches of a local database of 375 publicly available fungal genomes identified 2396 gene clusters containing homologs of at least three candidate cathinone genes, but none contained a transaminase, which is predicted to be required for cathinone synthesis (***Figure 6– figure supplement 2)***. We also found 309 clusters containing homologs of at least three candidate psilocybin genes, 175 of which contain a methyltransferase, which catalyzes the formation of psilocybin (***Figure 7–figure supplement 1; Table S7***). One notable locus in *Conidiobolus coronatus* (Entomophthorales) contained four aromatic-L-amino-acid decarboxylases and one methyltransferase (both functions are required for psilocybin biosynthesis), in addition to one helix-loop-helix transcription factor, one inositol phosphatase, and various amino acid, sugar and inorganic ion transporters (***Table S8***).

To test for psilocybin biosynthesis that could result from this single locus uncovered in *Conidiobolus coronatus* (Entomophthorales), eight representative *C. coronatus* isolates from a diverse set of hosts (***Table S9***) were acquired from the Agricultural Research Service Collection of Entomopathogenic Fungal Cultures (ARSEF). The eight isolates were used in both live-plating (direct exposure to spores) and injection entomopathogenicity assays on *Galleria mellonella*, a model insect for entomopathogenicity studies (***Panaccione and Arnold, 2017***). By 24-hours post-inoculation, all fungus-inoculated individuals were dead and by 48-hours all live plated individuals were symptomatic and many had died. Symptomatic individuals were collected at 48-hour post-inoculation and immediately placed at −80**°**C prior to maceration to permit screening of psilocybin and psilocin from both infected larvae and pure cultures of each isolate using a high resolution, accurate mass Orbitrap LC-MS. Neither compound was detected in macerated infected larvae or week-old fungal cultures of these same isolates on Sabouraud Dextrose Yeast Agar. However, *C. coronatus* isolate ARSEF 8715 (NRRL 28638), from which the genome and candidate locus was obtained, was not available and therefore was not included in the study.

Since genes encoding the synthesis of specialized secondary metabolites are occasionally inherited through non-vertical means, we assessed the potential for horizontal gene transfer of cathinone and psilocybin biosynthesis genes by constructing maximum likelihood phylogenetic trees for 22 high quality candidate pathway genes from *Mas. cicadina* and *Mas.* aff. *levispora* (NM) (***Figure 6–figure supplement 4; Figure 7–figure supplement 3; Table S10-S11***). High quality candidate genes were defined as sharing putative regulatory motifs in their 5’ upstream non-coding regions with either a transaminase or methyltransferase gene (which encode enzymes that produce cathinone or psilocybin, respectively), or homologous to genes in the cluster of interest in *C. coronatus*. All gene trees of these candidates had topologies consistent with vertical inheritance among the Entomophthorales with highly similar homologs present in other Entomophthoralean fungi. The phylogenies of candidate psilocybin pathway sequences gleaned from the preliminary metagenomic data are consistent with an origin of psilocybin biosynthesis independent from mushroom-forming Basidiomycetes, but other evolutionary origins cannot be ruled out until the exact genes underlying this pathway are confirmed.

### Phylogenomic resolution of Entomophthorales fungi and their behavior-modifying lifestyles

The metabolomics-based discovery of cathinone and psilocybin production in *Massospora* raises interesting questions about the broader evolutionary history of behavior-modifying traits across the Entomophthorales. In order to model evolution of such traits and establish whether AHT behavior seen in *Massospora* and other close relatives, including the only other known AHT-exclusive genus *Strongwellsea* (***Batko and Weiser, 1965; Roy et al., 2006***), reflects a single origin or multiple independent origin events, we conducted a phylogenetic analysis of a genome-scale data set of > 200 conserved orthologous proteins deduced from the *Mas. cicadina* and *Mas.* aff. *levispora* (NM) metagenomes along with 7 other representative taxa (***Figure 8–figure supplement 1***). These taxa span 6 of 20 described Entomophthorales genera and 3 of 4 genera with known active host transmission (***Hodge et al., 2017; Humber, 2012; Roy et al., 2006***).

**Figure 8.**
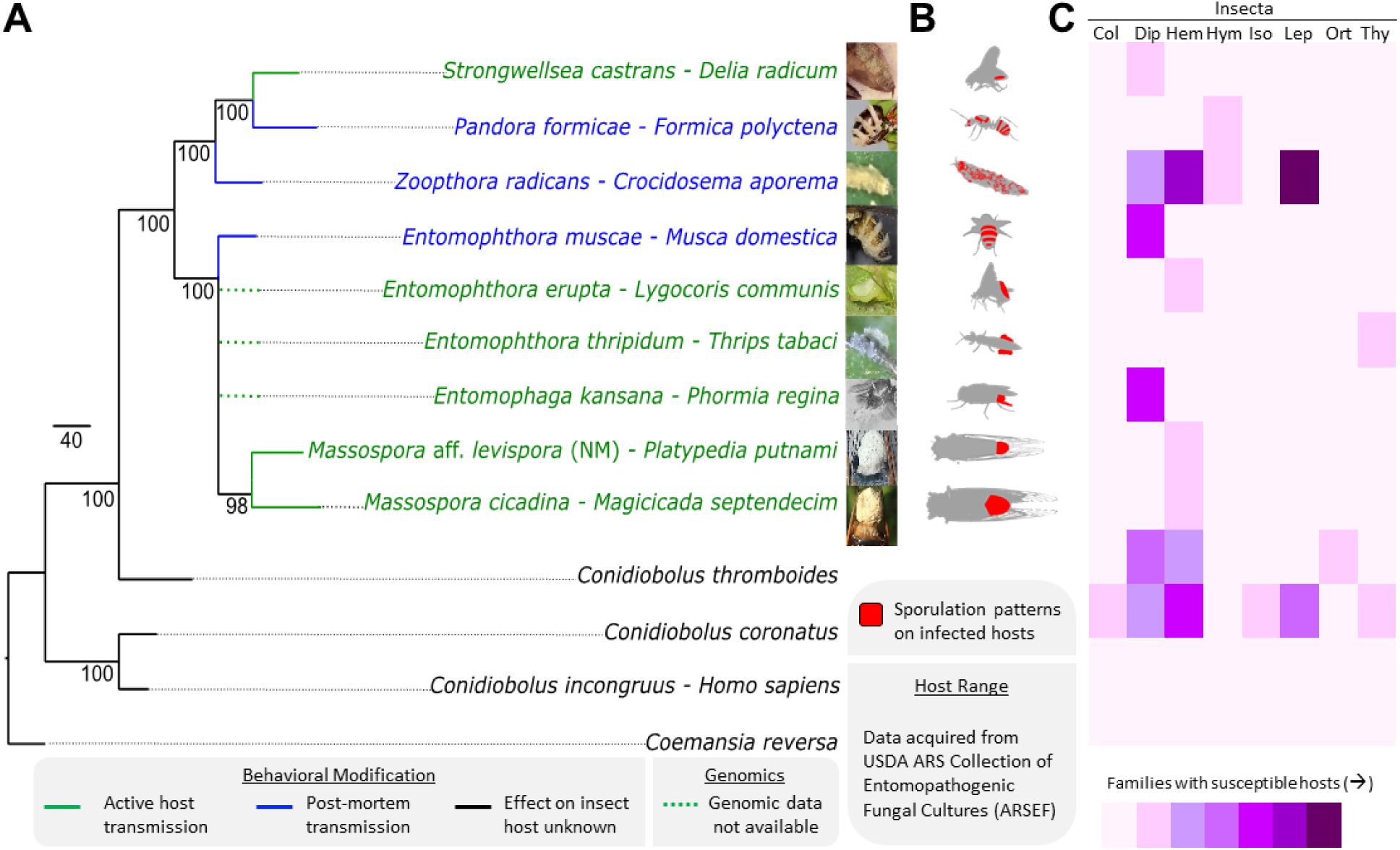
Phylogenomic relationships among behavior-modifying Entomophthoralean fungi, the behavioral and morphological host modifications they impose, and their host specificity. (A) RAxML phylogenetic tree of Entomopthoralean fungi (Zoopagomycota) based on the concatenated alignment of >200 conserved orthologous protein sequences with overlaid behavioral modification in infected hosts. Actual number of protein sequences per taxon denoted in gray. (B) External sporulation patterns, (C) Heat map showing both the host range within and across select insect orders: Col: Coleoptera, Dip: Diptera, Hem: Hemiptera, Hym: Hymenoptera, Iso: Isoptera, Lep: Lepidoptera, Ort: Orthoptera, and Thy: Thysanoptera. The species *Z. radicans* and *C. thromboides* may represent broader species complexes, of which each phylogenetic species may have a narrower host spectrum. Photos used with permission from co-authors as well as Florian Freimoser, Joanna Malagocka, Judith Pell, and Ruth Ahlburg.

Mapping of behavioral traits and external sporulation patterns onto the inferred phylogeny from the concatenated genome-wide alignment revealed three possible parsimonious reconstructions of AHT in Entomophthorales fungi (***Figure 8***). These topologies include either two independent gains of AHT, one of which is followed by a single loss, or a single gain in the most recent shared ancestor of *Massospora, Entomophthora*, and *Strongwellsea* followed by three losses, the latter of which suggests this behavior modification may have been the key innovation that led to the diversification of the Entomophthorales. At the same time, AHT entomopathogens exhibit a high degree of host specificity compared to their summit disease pathogens (***Figure 8–figure supplement 2***).

Follow-up metabolomics studies of other AHT fungi will likely help resolve these relationships especially if AHT was independently acquired by *Strongwellsea*.

## Discussion

One of the least explored frontiers in host-parasite interactions is the molecular basis for behavioral modification of hosts by fungal pathogens that promotes disease transmission. Recent studies of entomopathogenic *Ophiocordyceps* (Ascomycota) have identified neurogenic metabolites and enterotoxins that are possibly contributing to host manipulation (***De Bekker et al., 2017; Hughes, 2013***). Here, we report the surprising discovery that two species of *Massospora*, a genus of obligate entomopathogenic fungi infecting cicadas, produce psychoactive compounds during infection. These compounds are cathinone, an amphetamine only previously known from plants, and psilocybin, a tryptamine only previously known from Basidiomycete mushrooms. Prior work has demonstrated that *Massospora* can modify host behavior and we provide evidence for a chemical basis for this manipulation. This is the first evidence of amphetamine production in fungi and of psilocybin production outside the Basidiomycota.

Without exposure studies involving healthy cicadas (nymphs and adults), the mechanism(s) by which monoamine alkaloids modify cicada behaviors cannot be fully resolved. However, previous arthropod exposure studies have provided extensive insight into the effects of amphetamine and psilocybin on various behaviors (***Brookhart et al., 1987; Frischknecht and Waser, 1978; Witt, 1971***). For example, low doses (1-12 µg) of amphetamine similar to the cathinone levels detected in *Mas. cicadina* increased intraspecific aggression in ants (***Frischknecht and Waser, 1978***) and significantly depleted biogenic amines, altering feeding behavior in adult blow flies (*Phormia regina*) by decreasing responsiveness to weak gustatory stimuli (***Brookhart et al., 1987***). Psilocybin ingestion had less observable impact on behavior compared to amphetamine for the limited number of arthropods observed (***Witt, 1971***), but still may antagonize certain mycophagous insects that feed broadly on dung-loving basidiomycetes, thus conferring an evolutionary advantage to psilocybin-producing mushrooms (***Reynolds et al., 2018***). Psilocybin may also confer protection against predation, competition, and/or parasitism for a select few insects that exhibit indifference to psilocybin. For example, the dark-winged fungus gnat (Sciaridae), can successfully complete its lifecycle in fruiting bodies of psilocybin-containing *Psilocybe cyanescens* (***Awan et al., 2018***). Likewise, leafcutter ants (*Acromyrmex lobicornis*) have been observed actively foraging on *Psilocybe coprophila* fruiting bodies in Argentina, transporting basidiocarps back into the nest, possibly for defense purposes (***Masiulionis et al., 2013***). Although the effect of psilocybin on *Mas. levispora* (MI) differs somewhat in that the production of these compounds seem to benefit the parasite more than the host, the psilocybin may offer the insect some protection against attacks by the parasitoid fly *Emblemasoma auditrix* (Diptera: Sarcophagidae) (***Soper et al., 1976***).

While these secondary metabolites from *Massospora*, especially cathinone, may help improve endurance required to engage with other conspecifics given their debilitating infections, activity-level effects of amphetamine and psilocybin cannot easily explain the display of female-typical courtship behaviors by male cicadas (***Cooley et al., 2018***), except possibly in *P. putnami* (***Murphy and Redden, 2003***). Infected *P. putnami* males were found to produce significantly fewer crepitations (wing tapping behaviors required for mating) per observation period (***Murphy and Redden, 2003***). Fewer crepitations are associated with female-typical behavior, and the manifestation of these behaviors in males may indicate impaired temporal processing, a known effect of psilocybin in vertebrates (***Wittmann et al., 2007***). However, the atypical mating behaviors of other cicada species may be better explained by *Massospora*-mediated hormonal changes incidental to amphetamine and psilocybin production. For example, 2-hydroxyestradiol, a catechol estrogen and metabolite of estradiol and 3-O-acetylecdysone 2-phosphate, a metabolite of ecdysone, a steroidal prohormone of the major insect molting hormone 20-hydroxyecdysone, were both detected based on *m/z* and found to be upregulated in male *Platypedia putnami* with *Mas*. aff. *levispora* (NM)-infections compared to a mixed sex group *Magicicada septendecim* with *Mas. cicadina*-infections and healthy controls using global metabolomics (data not shown). These hormones may accumulate as a result of disruption of typical pathways (i.e. castration of testis) or they may be fungus-derived. Comparisons of *Mas.* aff. *levispora* (NM) fragmentation patterns with experimentally observed analytical standards are needed to validate these preliminary findings. Nevertheless, the differentially expressed “hypersexualization” between conidial- and resting spore-infected cicadas seems to support the latter hypothesis since changes in male behaviors cannot be incidental effects of castration or infection, or the observed behaviors would be seen in hosts accumulating either spore type. The basis of hypersexualization remains unknown; the compounds we identified may be involved wholly or partly in this phenomenon, and/or may serve other roles outside this observed phenotype.

Culturable fungi have yielded many antimicrobial drugs, medications, and industrial enzymes over the last century. The discovery of amphetamine production in fungi as well psilocybin production in the Zoopagomycota is consistent with the breadth of diverse secondary metabolite synthesis pathways that have evolved in fungi. However, the discovery of these alkaloids with limited genetic evidence of a common biosynthetic pathway raises questions about their proposed enzymatic route. The lack of evidence of a fungal kinase, an enzyme class which catalyzes the phosphotransfer step in psilocybin biosynthesis and of phenylalanine ammonia lyase (PAL), an enzyme class that catalyzes a reaction converting L-phenylalanine to ammonia and trans-cinnamic acid in the early steps of cathinone biosynthesis, coupled with the lack of many of these same pathway metabolites seems to support novel or altered biosynthetic pathways for both compounds.

Convergent pathways for de novo metabolite synthesis are known from fungi, such as pyridoxine biosynthesis in the phytopathogenic fungus *Cercospora nicotianae* (***Ehrenshaft et al., 1999***). Likewise, psychoactive cannabinoids, well known from *Cannabis sativa*, have been recently confirmed from liverwort, providing a marked example of convergent evolution of bioactive plant secondary metabolites (***Chicca et al. 2018***). Similarly, β-carboline alkaloids, are found in both plants and fungi, including in the Entomophthoralean fungus, *Conidiobolus coronatus*, where it is thought to interfere with insect serotonin levels and larval development (***Wrońska et al., 2018***).

The discovery of cathinone in *Massospora*-infected cicadas also raises interesting questions about the biological origin of amphetamine production in *Catha* plants and the possibility of cryptic fungal partners possibly related to *Massospora* involved in its biosynthesis. Over the last few decades, fungal endophytes have been identified as the source of ergot alkaloid production in fescue *(Festuca* spp.) and morning glories (*Ipomoea* spp.) (***Kucht et al., 2004; Lyons et al., 1986***). Culture-based studies of *Catha edulis* (***Alhubaishi and Abdel-Kader, 1991; Mahhoud, 2000***) and ephedrine-producing *Ephedra nebrodensis* (***Peláez et al., 1998***), have uncovered a diverse assortment of fungi, none of which have been screened for cathinone production. Interestingly, at least two cicada genera have been reported feeding on *Ephedra* plants (***Sanborn and Phillips, 2011; Wang et al., 2017***), which may indicate a tolerance to these compounds that are known to be toxic to other insects including bees (***Detzel and Wink, 1993***).

The results of this study provide a strong impetus for enhanced screening of Ephedraceous and Celastraceous plants for alkaloid-producing endophytes that may also have entomopathogenic capabilities. Interestingly, several recent studies have elucidated such tripartite relationships between insects, entomopathogenic fungi with known endophytic capability, and plant hosts whereby plants supply fungi with photosynthate in exchange for insect-derived nitrogen (***Barelli et al., 2015; Behie et al., 2012; Behie et al., 2017***).

Similarly, despite the very recent discovery of Hypocrealean fungal symbionts from the fat bodies surrounding the bacteriomes and/or the well-developed surface sheath cells of more than a dozen cicada species native to Japan (***Matsuura et al., 2018***), our results do not support the presence of cryptic domesticated monoamine alkaloid-producing fungal symbionts in the posterior region of healthy or outwardly asymptomatic wing-banger and periodical cicadas. Further studies should focus on resolving whether such domesticated symbionts are present in any North American cicada and how these fungi, if present, may interact with obligate pathogens such as *Massospora.*

The obligate lifestyle and ephemeral nature of many Entomophthoralean fungi have long constrained our understanding of these early diverging entomopathogens despite up to a century of observational studies into their behavior-modifying capabilities. We anticipate these discoveries will foster a renewed interest in early diverging fungi and their pharmacologically important secondary metabolites, which may serve as the next frontier for novel drug discovery and pave the way for a systems biology approach for obligate fungal pathogens spanning the known diversity of Kingdom Fungi.

## Methods

### Sampling of healthy and fungus-infected cicadas

Sampling of freshly infected and asymptomatic wing-banger cicadas and Brood VII periodical cicadas was carried out on private lands in CA (2018), NM (2017), and NY (2018) with permission from individual landowners. Infected Say’s cicada and Brood V periodical cicadas were collected from university-owned properties in MI (1998) and WV (2016), respectively. Collection of infected and healthy cicadas from all locations did not require permits. In CA, cicadas were collected under Exemption 5 of California Code of Regulations Section 650, Title 14 (https://nrm.dfg.ca.gov/FileHandler.ashx?DocumentID=157389&inline).

From May 28 to June 6, 2016, newly emerged Brood V 17-year periodical cicadas were visually inspected for *Massospora* fungal infections on the campus of West Virginia University in Morgantown, West Virginia, USA. Individuals with visible fungal plugs were transferred individually to sterile 20 mL vials, transported to the laboratory, and stored in a −20 C freezer until processing. Samples were obtained from 95 infected cicadas, of which 47 were from *Mag. cassini*, 41 from *Magicicada septendecim*, and seven from *Mag. septendecula*. A similar number of males and females were sampled across the three Brood V species, with 53 from females and 50 from males.

On June 1, 2016, additional geographically separated Brood V cicada populations were visually inspected for fungal infections. Seven infected specimens of *Mag. septendecim* and one of *Mag. cassini* were recovered from Hocking Co., OH. Concurrently, a co-author provided an additional 34 infected Brood V specimens of *Mag. septendecim* collected from across five OH counties including Ashland, Cuyahoga, Hocking, Licking, and Lorain counties. Eight female, *Massospora*-infected, Brood VI specimens of *Magicicada septendecim* were collected in June, 2017 from North Carolina. All samples were collected and stored as previously described.

Twenty-three *Mas. cicadina*-infected specimens of *Magicicada* collected from 1978-2002 were provided from the University of Connecticut, where they had been stored at room temperature, dried and pinned, since their collection. A second collection from Mount St. Joseph University provided fungal plugs from seven *Mas. cicadina*-infected specimens of *Magicicada* collected from 1989-2008, three of which were 13-year cicadas and four were 17-year cicadas.

In addition to periodic cicadas, two species of annual cicadas also were sampled and included in this study. On June 4, 2017, newly emerged banger-wing cicadas (*Platypedia putnami*) were inspected for *Massospora* infections 12 km northeast of Gallina, NM along the Continental Divide Trail. Sixteen infected individuals and one outwardly asymptomatic individual (to serve as a negative control) were identified and sampled as described above.

Nine *Mas. levispora*-infected specimens of *Okanagana rimosa* were collected by John Cooley from the University of Michigan Biological Station, Cheboygan and Emmet Counties, MI on June 20, 1998 (***Cooley et al., 2018; Cooley, 1999***). Infected individuals as well as outwardly asymptomatic individuals were placed immediately in individual 20-ml screw-cap vials containing 95% ethanol. Specimens were completely submerged and stored at 4 C until October 2017, when specimens were sent to WVU for processing.

Two independent populations of healthy cicadas, wing-banger cicadas from Trinity County, CA, USA and Brood VII periodical cicadas (*Mag. septendecim*) from Onondaga Hill, NY were sampled in early June, 2018 to serve as negative controls for targeted metabolomics. Five males and five female cicadas were collected for each cicada species, transferred individually to sterile 25-50 mL vials, and stored in a −80 C freezer until processing.

Cicadas were processed aseptically, with half of the visible fungal plug removed with a sterile scalpel and transferred to a 1.5 ml microcentrifuge tube for DNA extraction. Controls were processed similarly, where half of the most posterior segments as well as underlying host tissues were sampled. A majority of the remaining plug from infected individuals were used for global metabolomics analyses. Dry plugs were transferred aseptically to sterile 1.5 mL microcentrifuge tubes and stored at −80 C. A small sampling of spores was retained for morphological studies, although not all specimens were examined.

### Morphological studies of *Massospora* spp

Given the scarcity of *Massospora* sequences from NCBI GenBank, morphological comparisons were necessary to determine if data on putatively assigned *Massospora* species agreed with that of previously reported measurements for conidial and resting spore stages (***Soper, 1963; Soper, 1974***). A portion of select fungal spore masses was harvested with a sterile scalpel and mounted in lactophenol for examination with light field microscopy. Slip covers were fastened with nail polish to allow slides to be archived and reexamined when necessary. Mean conidial/resting spore length x width and cicada host provided the main criteria to differentiate *Massospora* species. Slides were examined and photographed using a Nikon Eclipse E600 compound microscope (Nikon Instruments, Melville, New York) equipped with a Nikon Digital Sight DS-Ri1 high-resolution microscope camera. A sampling of 25 spores from each slide mount were measured using Nikon NIS-Elements BR3.2 imaging software. A total of 51 *Massospora*-infected cicadas were examined including 26 *Mas. cicadina* (15 conidial and 11 resting spore isolates), nine *Mas. levispora* (eight conidial and one resting spore isolates), and 15 *Mas. platypediae* (14 conidial isolates and one resting spore isolate).

### DNA extraction, amplification, sequencing, and assembly

Fungal genomic DNA was extracted from harvested fungal plugs using a modified Wizard kit (Promega, Madison, WI, USA) as previously described (***Short et al., 2015***). DNA was suspended in 75 µl Tris-EDTA (TE) buffer and stored at −20 C. Portions of the nuclear 18S (SSU) and 28S (LSU) ribosomal DNA regions were amplified with the primer combinations NS24/NSSU1088R (***Hodge et al., 2017***) and LR0R/LR5 (***Vilgays and Hester, 1990***) using BioLine PCR kits (Bioline USA Inc.,Taunton, MA, USA). EFL, an EF-like gene found in some groups of early diverging fungi including the Entomophthorales, was amplified using primers EF1-983F and EF1aZ-1R (***James et al., 2006***). Eight degenerate primers pairs (***Table S1***), three of which were validated against a positive control, were designed to target three psilocybin biosynthesis enzymes, PsiD (l-tryptophan decarboxylase), PsiK (4-hydroxytryptamine kinase), and PsiM (norbaeocystin methyltransferase), described from hallucinogenic mushrooms, *Psilocybe cubensis* and *Psilocybe cyanescens* (***Fricke et al., 2017, Reynolds et al., 2018***), were used to attempt to amplify *Psilocybe* psilocybin biosynthesis genes in *Mas. platypediae.*

PCR conditions were as follows. LSU: initial denaturation at 94°C for 2 minutes, 35 cycles of denaturation at 94°C for 30 seconds, annealing at 51.1°C for 45 seconds, and elongation at 72°C for 90 seconds, and final elongation at 72°C for 5 minutes; SSU: initial denaturation at 94°C for 5 minutes, 36 cycles of denaturation at 94°C for 30 seconds, annealing at 49°C for 30 seconds, and elongation at 72°C for 60 seconds, and final elongation at 72°C for 7 minutes. EFL: initial denaturation at 94°C for 5 minutes, 30 cycles of denaturation at 95°C for 60 seconds, annealing at 54°C for 60 seconds, and elongation at 72°C for 90 seconds, and final elongation at 72°C for 7 minutes. PsiD: initial denaturation at 95°C for 4 minutes, 30 cycles of denaturation at 95°C for 45 seconds, annealing at 61°C for 45 seconds, and elongation at 72°C for 45 seconds, and final elongation at 72°C for 7 minutes. PsiK: initial denaturation at 95°C for 5 minutes, 35 cycles of denaturation at 95°C for 45 seconds, annealing at 52.5°C for 50 seconds, and elongation at 72°C for 50 seconds, and final elongation at 72°C for 5 minutes. PsiM: initial denaturation at 95°C for 5 minutes, 35 cycles of denaturation at 95°C for 45 seconds, annealing at 54°C for 50 seconds, and elongation at 72°C for 40 seconds, and final elongation at 72°C for 5 minutes.

Amplified products were visualized with SYBR gold (Invitrogen, Grand Island, NY, USA), and then were loaded onto a 1.0-1.5%, wt/vol agarose gel made with 0.5-1% Tris-borate-EDTA buffer (Amresco, Solon, OH, USA). Psilocybin biosynthesis (Psi) products were cleaned with Zymo Research (Irvine, CA, USA) DNA Clean and Concentrator-5 kit. All products were purified with ExoSAP-IT (Affymetrix, Inc., Santa Clara, CA, USA) and submitted to Eurofins MWG Operon (Huntsville, AL, USA) for Sanger sequencing. Chromatograms were assembled and inspected using Codon Code aligner and phylogentic analyses conducted using MEGA 7 (***Kumar et al., 2016***).

The *Mas. platypediae* (syn. *Mas.* aff. *levispora* (NM)) metagenome assembly was generated from 390 Million 2×150 bp raw reads. Reads were quality trimmed with sickle using default parameters (***Joshi and Fass, 2011***). Metagenome assembly was performed with megahit with “--presets meta-sensitive” option and utilizing GPU cores for accelerated matching speed to produce a genome assembly of ∼3 Gb (2.1 million contigs; N50= 2784 bp). The *Mas. cicadina* assembly was generated with SPAdes using default parameters. Both assemblies were further cleaned for vector and primer contamination with NCBI vecscreen protocol (https://www.ncbi.nlm.nih.gov/tools/vecscreen/) and implemented in package (https://github.com/hyphaltip/autovectorscreen). Several rounds of BLASTN based screening were applied to remove contaminated contigs or regions. Further screening of contigs were performed during genome submission process to NCBI to remove highly similar matches to plant or insect sequences. These Whole Genome Shotgun projects have been deposited at DDBJ/ENA/GenBank under the accessions QKRY00000000 (*Mas. platypediae*) and QMCF00000000 (*Mas. cicadina*). The versions described in this paper are versions QKRY01000000, QMCF01000000.

### Phylogenetic analysis of *Massospora* spp

Phylogenetic trees were constructed using LSU, SSU, and EFL sequences generated from 22, 26, and 20 *Massospora* sequences, respectively, representing three *Massospora* species, *Mas. cicadina, Mas. levispora* and *Mas. platypediae*. The concatenated set (LSU+SSU) consisted of four sequences, two from *Mas. cicadina* and one each from *Mas. levispora* and *Mas. platypediae*. Nine additional published sequences spanning the known diversity of the Entomophthorales were included to help resolve relationships among *Massospora* and its close allies (***Hodge et al., 2017***). One isolate of *Conidiobolus pumilus* (strain ARSEF 453) served as an outgroup. Sequences were aligned using CLUSTAL-W (***Larkin et al., 2007***). Alignments were visually inspected, after which optimized nucleotide substitution model (Tamura 3-parameter + G) was selected using ModelTest (***Posada and Crandall, 1998***). Partial deletion with site coverage cutoff of 95% was used. Maximum likelihood (ML) trees were estimated using MEGA 7. Phylogenetic support was assessed by bootstrap analysis with 1000 replicates using MEGA 7.

### Phylogenomic analyses of Entomophthoralean fungi

Multi-gene phylogenetic analysis of protein coding genes was also performed to assess confidence in the phylogenetic relationships. Since the metagenome assemblies were highly fragmented due to high repetitive sequence content, a complete genome annotation was not undertaken. Instead a targeted similarity search identified contigs encoding homologs of a collection of generally single copy protein coding genes developed for the 1000 Fungal genomes project. These 434 “JGI_1086” markers are available as Hidden Markov Models (https://github.com/1KFG/Phylogenomics_HMMs). Consensus sequences generated from the JGI_1086 markers using hmmemit (HMMER v3) (***Eddy, 1998***) were used as queries in a translated search of the *Massospora* metagenome assemblies with TFASTY (***Pearson et al., 1997***) using a permissive expectation cutoff value of ‘0.001’. These candidate contigs were then annotated with funannotate pipeline (https://github.com/nextgenusfs/funannotate) with informant protein sequences of a collection Zoopagomycota fungi, *de novo* gene prediction tools Augustus (***Stanke and Waack, 2003***) and GeneMark ES+ET (***Lomsadze et al., 2005***).

The predicted proteins for the targeted *Massospora* contigs were analyzed in the PHYling pipeline with proteins from the annotated genomes of the Kickxellomycotina fungus *Coemansia reversa* NRRL 1564, the Entomophthoromycotina fungi, *Conidiobolus coronatus* NRRL 28638, *Conidiobolus incongruus* B7586, *Conidiobolus thromboides* FSU 785) and the predicted ORFs from assembled transcriptomes of Entomophthoromycotina *Entomophthora muscae, Pandora formicae, Strongwellsea castrans, Zoophthora radicans* ATCC 208865. Briefly PHYling (https://github.com/stajichlab/PHYling_unified; DOI: 10.5281/zenodo.1257002) uses a preselected marker set of HMMs (https://github.com/1KFG/Phylogenomics_HMMs; DOI: 10.5281/zenodo.1251477) to identify best homologous sequences, aligns, trims, and builds a concatenated phylogenetic tree or individual gene trees. Searches of the annotated genomes of most species identified between 397 (*C. coronatus*) and 427 (*S. castrans*) of the 434 marker genes in each species. Searches of the *Mas. cicadina* and *Mas. platypediae* targeted annotation set identified 202 and 237 of the 434 marker sets respectively. This lower number indicates a less complete and highly fragmented metagenome assembly but these ∼200 protein coding genes provide sufficient phylogenetic information to reconstruct species relationships. The 10-taxon tree was constructed with IQ-TREE (v1.6.3; ***Hoang et al., 2017; Nguyen et al., 2014***) using the concatenated alignment and 433 partitions and 1000 ultrafast bootstrap support computation (-bb 1000 -st AA -t BIONJ -bspec GENESITE). All marker gene sequences, alignment and partition files are available at DOI: 10.5281/zenodo.1306939.

### Fungal metabolite extraction from *Massospora* plugs and healthy cicadas

Fungal plugs or healthy cicada abdomens were weighed and transferred into sterile 1.5 mL microcentifuge tubes. Metabolite extracts from each fungal sample were normalized to a final concentration of 10 mg of fungal tissue to 1 mL and polar metabolites were extracted in a buffer consisting of 80% HPLC grade methanol and 20% water. Samples then were sonicated for 10 minutes. The sonicated samples were centrifuged at 14,000 x g for 15 minutes at 4°C to pellet proteins and cellular debris. The supernatant was removed and dried down using lyophilization prior to analysis. Samples were reconstituted in 100 µL of 50% methanol and 50% water for mass spectrometry analysis.

Mycelia from cultures of *Conidiobolus coronatus* were harvested by scraping 7-day old cultures grown on ½ strength Sabouraud dextrose agar amended yeast extract. Whole *Con. coronatus-*infected greater wax moth larvae (*Galleria mellonella*) were harvested at 48 hours post inoculation. Both tissues were macerated in a buffer as previously described.

### Analytical procedure and LC-MS conditions for metabolomics

The Agilent LC-MS system (Agilent Technologies, Santa Clara, CA, USA) included an Agilent 1290 Infinity quaternary ultra-high pressure liquid chromatography (UHPLC) pump. This LC system was coupled to an Agilent 6530 quadrupole time of flight (Q-ToF) mass spectrometer with electrospray ionization. The chromatographic separations were performed using a Merck polymeric bead based ZIC-pHILIC column (100mmx2.1mm, 5µm).

Mobile phase A consisted of 10 mM ammonium acetate and mobile phase B consisted of 100% acetonitrile. Metabolites were separated on a gradient that went from 95% mobile phase B to 5% mobile phase B over 20 minutes with 7-minute re-equilibration at the end of the chromatographic run.

The mass spectrometer was operated alternately in positive and negative ion mode for the analyses. Samples for global metabolomics consisted of fungal plug extracts from 5 *Mas. cicadina*-infected Brood V periodical cicadas, 5 *Mas. platypediae*-infected wing-banger cicadas from NM, and 4 technical reps of excised posterior segments from a single healthy brood V periodical cicada or a healthy *P. putnami* from NM. Data dependent MS/MS was performed on triplicate pooled samples for fragmentation and library searching. For *Mas. cicadina*, pooled samples consisted of 5 independently extracted Brood V periodical cicadas collected on the campus of WVU in 2016. For *Mas. platypediae* pooled sampled consisted of 5 independently extracted *P. putnami* annual cicadas collected in NM in 2017. Controls consisted of excised posterior segments from two healthy brood V periodical cicada or a healthy *P. putnami* from NM.

### Data processing, metabolite identification, and statistical data analysis for global metabolomics

The LC-MS/MS data acquired using Agilent Mass Hunter Workstation were processed in Agilent Profinder software version 2.3.1 for batch analysis. The datasets were subjected to spectral peak extraction with a minimum peak height of 600 counts and the charge state for each metabolite was restricted to two. Further, retention time and mass alignment corrections were performed using Agilent Profinder software version 2.3.1 on the runs to remove non-reproducible signals. The resulting features then were exported to Mass Profiler Professional (MPP) software version 2.4.3 (Agilent Technologies, Santa Clara, CA, USA) for statistical analysis. The extracted features were imported into MPP software for statistical analysis. Principle Component Analysis (PCA) was performed to check the quality of the samples after which the data containing filtered features were processed by one-way ANOVA to ascertain differences between control and fungal groups. Only the analytes with p values < 0.05 and fold change (FC) > 20 were treated as statistically significant. Additionally, multiple test corrections using Bonferroni were applied to reduce false positives and false negatives in the data.

### Targeted metabolomics

The absolute quantification of cathinone, psilocin and psilocybin from the *Mas. cicadina* and *Mas. platypediae* fungal plug metabolite extracts described above, *Mas. levispora* extracts that were stored long-term were not able to be analyzed for global metabolite changes were able to be analyzed in this targeted experiment. This analysis was performed using a 5500 QTRAP mass spectrometer (SCIEX Inc., Concord, Ontario, Canada). Chromatographic separations were performed using an HPLC system consisting of two Shimadzu LC 20 AD pumps that included a degasser and a Shimadzu SIL 20 AC auto sampler (Shimadzu Corp. Kyoto, Kyoto Prefecture, Japan). Analytes were separated on a 3.0×150 mm Imtakt Scherzo SW-C18 column (Imtakt USA, Portland, Oregon). The HPLC column oven was set to 40°C for the analysis, with an 8 µL injection volume for each sample. Mobile phase A consisted of HPLC grade H2O with 0.1% formic acid, while mobile B consisted of acetonitrile with 0.1% formic acid. The LC gradient started 5% B for 0.5 minutes and then increased to 95% B over 5 minutes followed by a re-equilibration to starting conditions for a period of 2.1 minutes. The flow rate was set to 0.3 mL/minute.

Targeted metabolites were quantified using multiple reaction monitoring. The transitions that were monitored for cathinone were *m/z* 150.1→106.3. The transitions that were monitored for psilocybin were *m/z* 284.5→205.2 and 240.1. Psilocin was monitored at *m/z* 205.2→116.3. Mass spectrometer source conditions were set to the following: curtain gas = 15, ion spray voltage= 4800 V, temperature= 600 °C, gas 1= 45, gas 2= 50. Calibration samples were prepared in 50 % methanol/H2O. Calibration concentrations ranged from 0.05 to 10 ng/mL of both psilocin and psilocybin and 1.25-25 ng/mL for cathinone. In order to assess variation in the assay duplicate standard curves were analyzed at the beginning and at the end of the analytical run. Absolute quantification data was processed using Analyst software ver. 1.6.3 (SCIEX Inc., Concord, Ontario, Canada).

### Targeted quantification of CoA dependent pathway intermediates

Targeted detection of CoA dependent pathway intermediates including trans-cinnamoyl-CoA, 3-hydroxy-3-phenylpropionyl-CoA, 3-oxy-3-phenylpropionyl-CoA, and benzoyl-CoA was performed using a 5500 QTRAP mass spectrometer (SCIEX Inc., Concord, Ontario, Canada). Chromatographic separations were performed using an HPLC system consisting of two Shimadzu LC 20 AD pumps that included a degasser and a Shimadzu SIL 20 AC auto sampler (Shimadzu Corp. Kyoto, Kyoto Prefecture, Japan). Analytes were separated on a 3.0 x 150 mm Imtakt Imtrada column (Imtakt USA, Portland, Oregon). Mobile phase A consisted of 10 mM ammonium acetate at pH 9.0 and mobile phase B consisted of 100% acetonitrile. CoA intermediates were separated on a gradient that went from 95% mobile phase B to 5% mobile phase B over 15 minutes with 3-minute re-equilibration at the end of the chromatographic run. Absolute quantification data was processed using Analyst software ver. 1.6.3 (SCIEX Inc., Concord, Ontario, Canada).

### Targeted screening of psilocybin and psilocin from *Conidiobolus coronatus* and 2018-collected healthy *Platypedia putnami* (CA) and cathinone from *Magicicada septendecim* (Brood VII)

Targeted cathinone, psilocybin and psilocin experiments were performed using an Accela 1250 UHPLC coupled to a Q Exactive Orbitrap Mass Spectrometer (Thermo Scientific, San Jose, CA) operating in positive ion mode. An Agilent Zorbax C18 column (2.1 mm diameter by 10cm length) was loaded with 2 µL of extracts. Separations were performed using a linear gradient ramping from 95% solvent A (water, 0.1% formic acid) to 90% solvent B (acetonitrile, 0.1% formic acid) over 6 minutes, flowing at 300 µL/min.

The mass spectrometer was operated in targeted-SIM/data-dependent MS2 acquisition mode. Precursor scans were acquired at 70,000 resolution with a 4 *m/z* window centered on 285.1005 *m/z*, 205.1341 *m/z*, and 150.0913 *m/z* (5e5 AGC target, 100 ms maximum injection time). Charge states 2 and higher were excluded for fragmentation and dynamic exclusion was set to 5.0 s.

### Search for candidate psilocybin and cathinone biosynthetic genes

We used multiple approaches to identify candidate psilocybin and cathinone biosynthetic genes (***Figure 6–figure supplement 1***). We first performed tBLASTn searches of *Massospora* metagenome assemblies using four characterized psilocybin biosynthesis proteins from *Psilocybe cubensis* (***Fricke et al., 2017***) (norbaeocystin methyltransferase (accession P0DPA9.1); L-tryptophan decarboxylase (accession P0DPA6.1); 4-hydroxytryptamine kinase (accession P0DPA8.1); cytochrome P450 monooxygenase (accession P0DPA7.1)) and all 42 candidate cathinone biosynthesis proteins identified in a transcriptomic study of *Catha edulis* (retrieved from S1 dataset and Table 2 of ***Groves et al., 2015***). We next subtracted aligned sequences between the two assemblies using NUCmer and PROmer from MUMMER v.4.0.0beta2 in order to identify sequence unique to each assembly (***Marçais et al., 2018***). Finally, we used profile HMM of the protein family (Pfam) domains found in characterized psilocybin and cathinone biosynthetic genes to search the 6-frame amino acid translations of the *Massospora* metagenomic assemblies with HMMER3 v.3.1b2 (***Eddy, 2011; Table S2***). We searched with all SAM and SAM-like methyltransferase domains because many are similar to the S-adenosyl-l-methionine (SAM)-dependent methyltransferase domain in the psilocybin pathway’s norbaeocystin methyltransferase. Unless stated otherwise, all BLAST searches were conducted using the BLAST suite v.2.2.25+ with a max. evalue threshold=1e-4 (***Altschul et al., 1990***) against a local database containing 375 publically available fungal genomes (***Table S5***).

All subsequent steps were performed on the set of sequences retrieved by profile HMM search. We filtered out bacterial, protist and insect sequences by aligning hits to NCBI’s non-redundant protein database (last accessed March 14^th^, 2018) using BLASTp. We annotated contigs containing filtered hits using WebAUGUSTUS with *C. coronatus* species parameters using the option “predict any number of (possibly partial) genes” and reporting genes on both strands (***Hoff and Stanke, 2013***). We manually verified the coordinates of CDS regions by using the hits as queries in a BLASTx search (min similarity=45%) to the local proteome database. The predicted amino acid sequences of all candidate genes, as well as the nucleotide sequences and coordinate files of annotated contigs are available at DOI: 10.5281/zenodo.1306939.

To investigate putative ungapped regulatory motifs associated with candidate gene sequences, we used a custom Perl script to extract up to 1500bp upstream of each gene’s start codon and submitted each set of extracted sequences to command line MEME v.4.12.0 (max evalue=5e-2; min width=25; max width=100; max motifs=10) (***Bailey and Elkan, 1994***). Motifs were manually inspected and those consisting solely of homopolymer repeats were discarded. Because the *Massospora* assemblies were themselves too fragmented to assess if candidate genes were found in clusters, we searched for evidence of clusters containing candidate gene homologs retrieved from the local fungal genome database using BLASTp and a custom Perl script (available at https://github.com/egluckthaler/cluster_retrieve). Homologs were considered clustered if separated by 6 or fewer intervening genes. Only clusters with at least 3 genes were retained. Genes at clustered loci of interest were annotated with eggNOG-mapper v.0.99.1 (***Huerta-Cepas et al., 2017***) based on orthology data from the fungal-specific fuNOG database (***Huerta-Cepas et al., 2016***).

We designated genes as “high-quality” candidates of interest if their upstream regions contained a motif also present in the upstream region of a predicted transaminase or methyltransferase gene. We used these criteria to help define candidates of interest because methyltransferase and transaminase enzymes catalyze the formation of psilocybin and cathinone, respectively.

To investigate the evolution of the 22 high quality candidate gene sequences (***Table S10***), we first divided sequences into groups based on the profile HMM that initially retrieved them. Sequences from each group were used as queries to retrieve hits from the local fungal genome database (***Table S5***) which was modified at this point to also include all candidate genes identified from both *Massospora* assemblies, as well as the metatranscriptomes of *Conidiobolus incongruus* B7586, *Entomophthora muscae* (ARSEF 13376), *Zoophthora radicans* ATCC 208865, *Strongwellsea* sp. (HHDFL040913-1), and *Pandora formicae* (***Malagocka et al., 2015***) using BLASTp.

The *Strongwellsea* sp. transcriptome was based on total RNA extracted from a single infected *Delia radicum* caught in an organically grown cabbage field in Denmark: Sørisgaard (55.823706, 12.171149). The fly had the characteristic *Strongwellsea-*caused open hole on the side of the abdomen and was snap-frozen in liquid nitrogen as quickly as possible. Illumina sequencing yielded ca. 20 million 150 bp paired-end reads that were assembled using Trinity 2.2.0 (***Grabherr et al., 2011***). Transcripts were filtered so that only the highest expressed isoform for each transcript and transcripts assigned to kingdom Fungi using MEGAN4 (***Huson et al., 2011***) were retained.

Each set of queries and associated hits (with two exceptions, see below) were then aligned with MAFFT v.7.149b using the --auto strategy (***Katoh and Standley, 2013***), trimmed with TrimAL v.1.4 using the –automated1 strategy (***Capella-Gutiérrez et al., 2009***), and had sequences composed of >70% gaps removed. Sets of sequences retrieved using candidates with the ‘p450’ or ‘aldo_ket_red’ Pfam HMMs were too large to construct initial alignments. Instead, all retrieved hits and queries were first sorted in homolog groups by submitting the results of an all vs. all BLASTp to OrthoMCL v.2.0 (inflation = 1.5) (***Li et al., 2003***). All sequences in the homolog groups containing the query candidate genes were then aligned and trimmed as above. Each trimmed alignment was then used to construct a phylogenetic tree with FASTTREE v.2.1.10 using default settings that was then midpoint rooted (***Price et al., 2010***). For each high quality query sequence present in the tree, a strongly supported clade containing the query sequence and closely related Entomophthoralean sequences was extracted. Sequences in the extracted clade were re-aligned and trimmed (as above) and used to build final maximum likelihood trees using RAxML v8.2.11 with 100 rapid bootstraps (***Stamatakis, 2014***; ***Figure 6– figure supplement 4; Figure 7–figure supplement 3***). Final RAxML trees were rooted based on the outgroup in the clade extracted from the midpoint rooted FASTTREE.

## Acknowledgments

Special thanks to the Drug Enforcement Administration (DEA) for guidance on discovery and handling of Schedule 1 controlled substances. Genomes and transcriptome sequencing of *C. thromboides, C. coronatus*, and *Z. radicans* were supported by the Department of Energy’s Joint Genomes Institute under the JGI Community sequencing program project 1978 ‘Genomics of the early diverging lineages of fungi and their transition to terrestrial, plant-based ecologies’ supported by the Office of Science off the US Department of Energy under contract DE-AC02-05CH11231.

## Funding

G.R.B. was supported by Protea Biosciences and NSF grant DBI-1349308, J.C.S. by NSF DEB 1638999, J.E.S. by NSF DEB 1441715, W.J.D. and T.Y.J. by NSF DEB 1441677, D.G.P by NIH 2R15GM114774-2, H.H.D.F.L. by the Villum Foundation (grant number 10122), T.G. by NSF grant CHE-1608149, J.R.C. and C.S. by NSF DEB 1655891, and M.T.K. by funds from West Virginia Agricultural and Forestry Experiment Station and NSF DBI-1349308.

## Author contributions

G.R.B., J.S., W.J.D., T.Y. J., D.G.P., and M.T.K. conceived of the study. J.R.C., C.S., A.M.M., M.C.B., G.K., K.T.H., R.A.H., T.G., D.M., and N.A. provided archived material and/or fresh specimens. G.R.B., A.M.M., M.C.B., K.L.W., C.M.S., E.J.S., A.M.M., T.K. and M.T.K. performed laboratory work with the help of D.P.G.S. and D.G.P. G.R.B., E.G.-T., J.C.S., J.S., W.J.D., T.Y.J., J.E., H.H.D.F.L., M.D.M., and M.T.K. analyzed data. G.R.B., E.G.-T., J.C.S., J.S., W.J.D., J.R.C., J.E., C.S., and M.T.K. wrote the manuscript with input from all coauthors;

## Competing interests

Authors declare no competing interests

## Data and materials availability

Whole Genome Shotgun projects have been deposited at DDBJ/ENA/GenBank under the accessions QKRY00000000 (*Mas. platypediae or Mas.* aff. *levispora* (NM)) and QMCF00000000 (*Mas. cicadina*). The versions described in this paper are QKRY01000000 and QMCF01000000. All phylogenomic marker and candidate cathinone and psilocybin gene sequences are available at DOI: 10.5281/zenodo.1306939.

## Supplementary Figures

**Figure 1–figure supplement 1.**
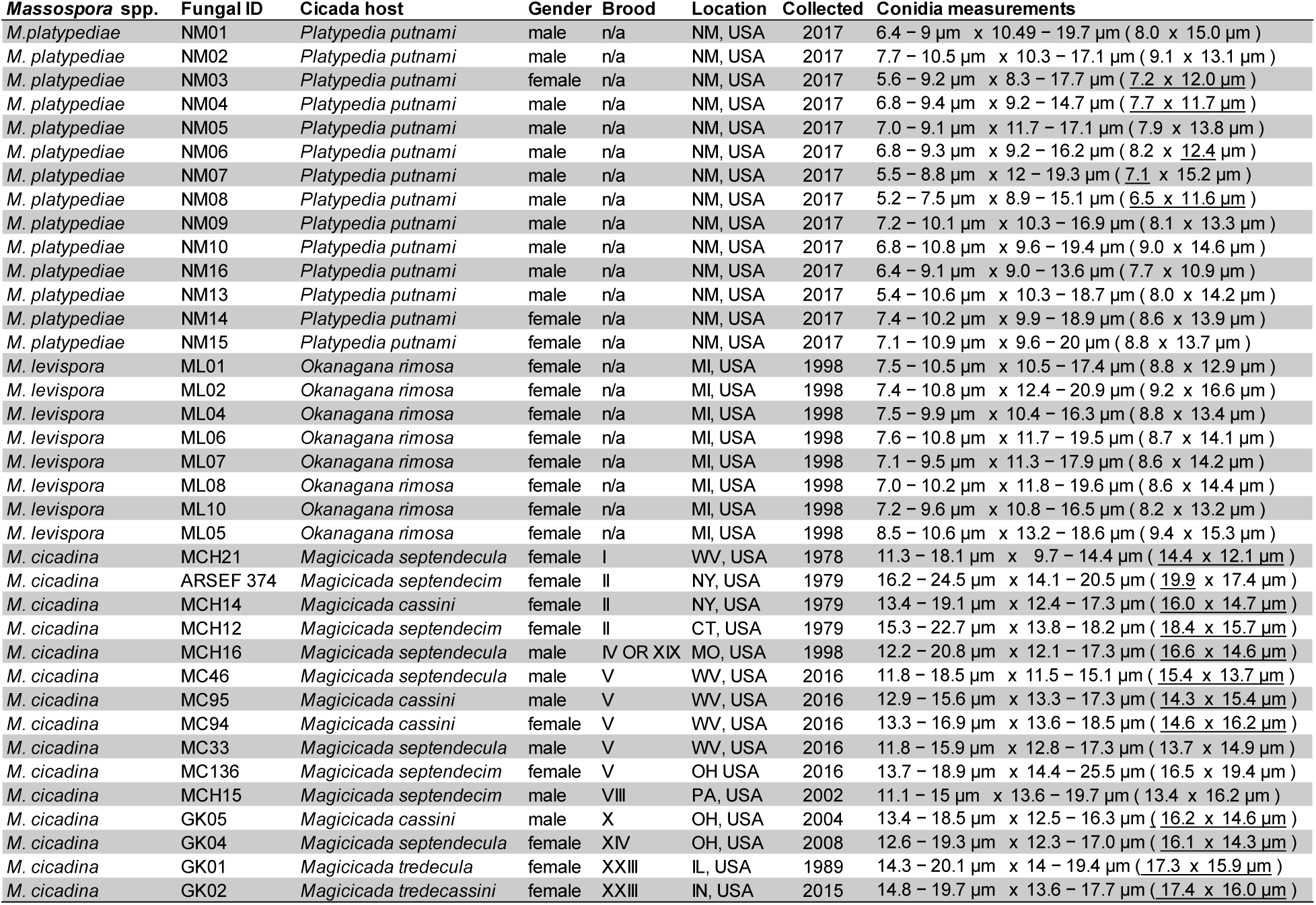
*Massospora* spp. conidial measurements based on the average of 25 measured spores per fungal isolate. Underlined mean values in parentheses denote that mean length and width fell within the previously reported ranges for that given species.

**Figure 1–figure supplement 2.**
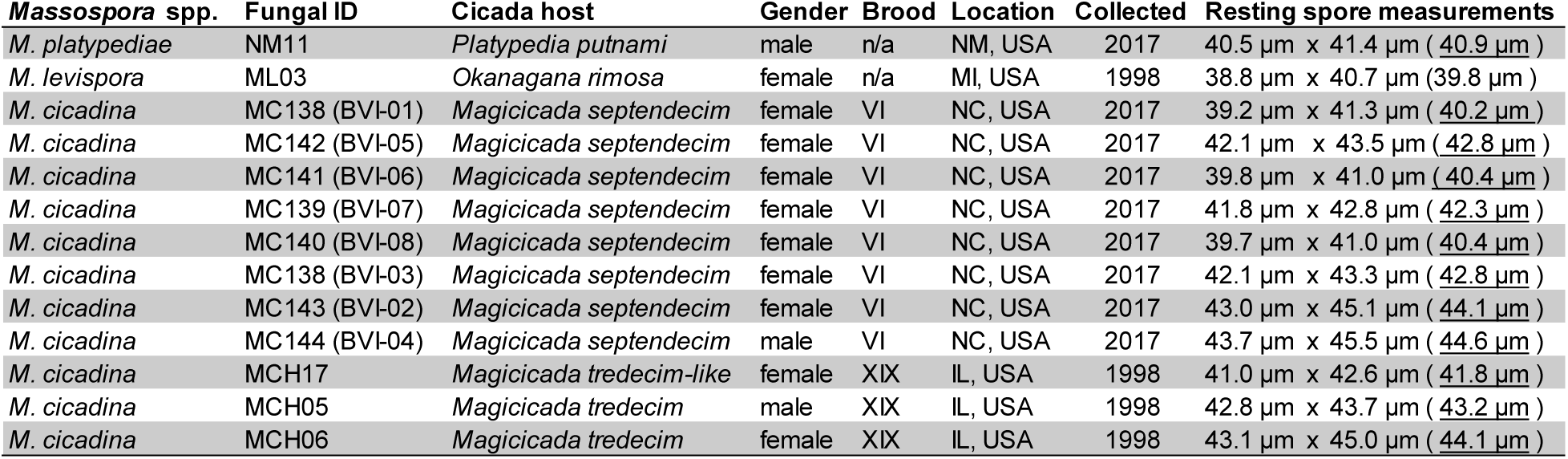
*Massospora* spp. resting spore measurements based on the average of 25 measured spores per fungal isolate for *Mas. cicadina* and 50 spores per isolate for *Mas. levispora* and *Mas. platypediae*. Underlined mean values in parentheses denote that mean length and width fell within the previously reported ranges for that given species.

**Figure 3–figure supplement 1.**
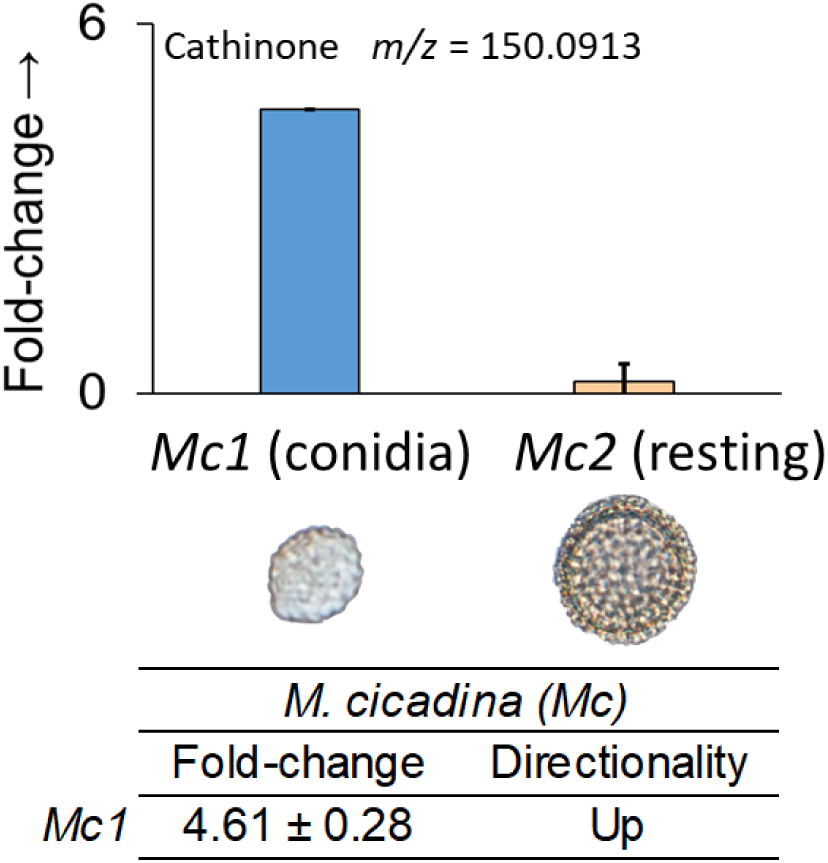
Global metabolomics fold-change comparisons of cathinone between *Mas. cicadina* conidal plugs (*Mc1*) and resting spore plugs (*Mc2*). Value is average of five biological replicates.

**Figure 5–figure supplement 1.**
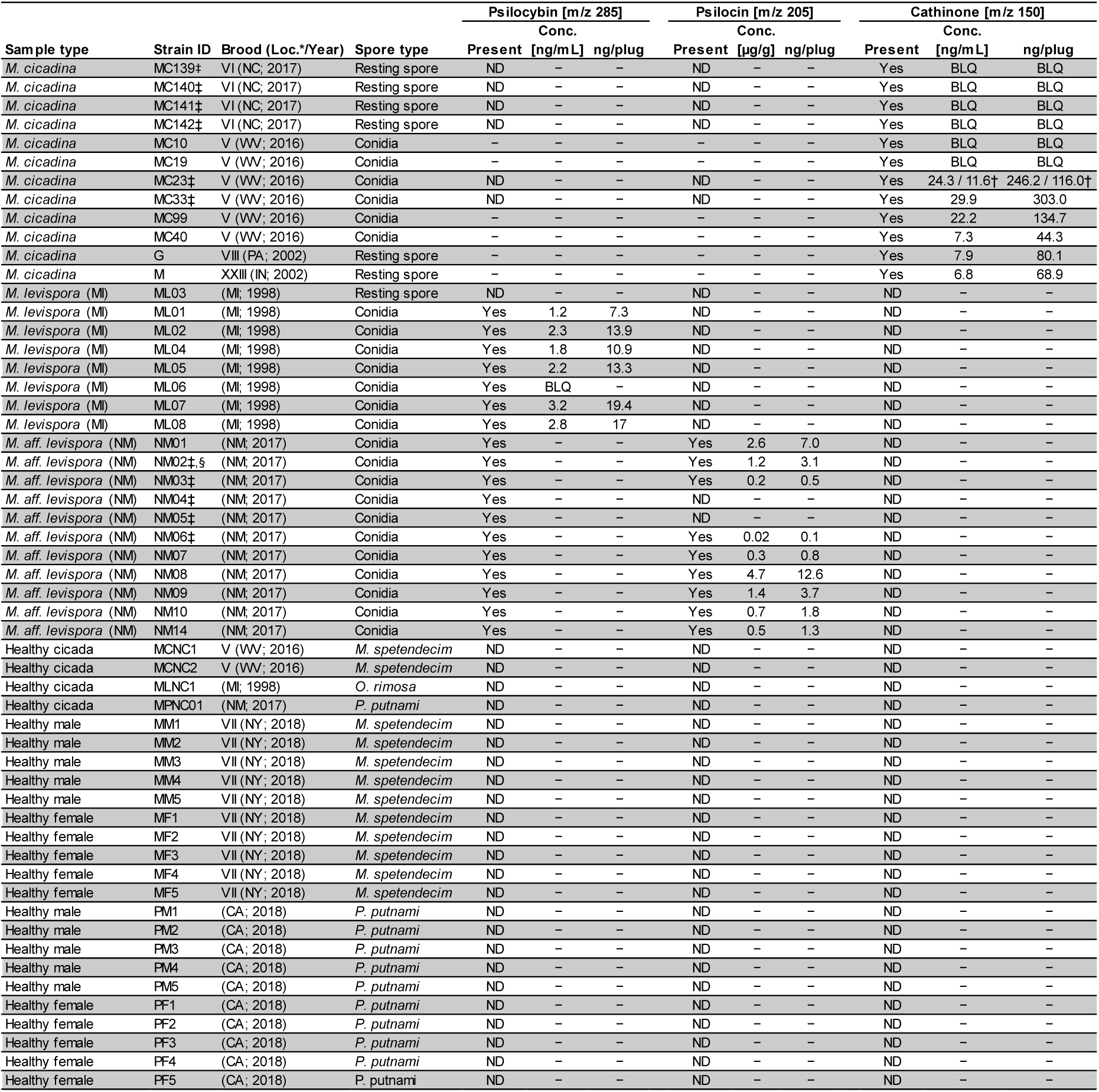
Isolate table for absolute quantification of psilocybin, psilocin, and cathinone by LC-MS/MS. Samples that fell outside the limit of quantification were scored as either present or absent only. ND, not detected; BLQ, below limit of quantification; * U.S. state abbreviation, † indicates samples were included in two independent runs, ‡ isolates used in global metabolomics studies, § isolate from which metagenome was sequenced.

**Figure 6–figure supplement 1.**
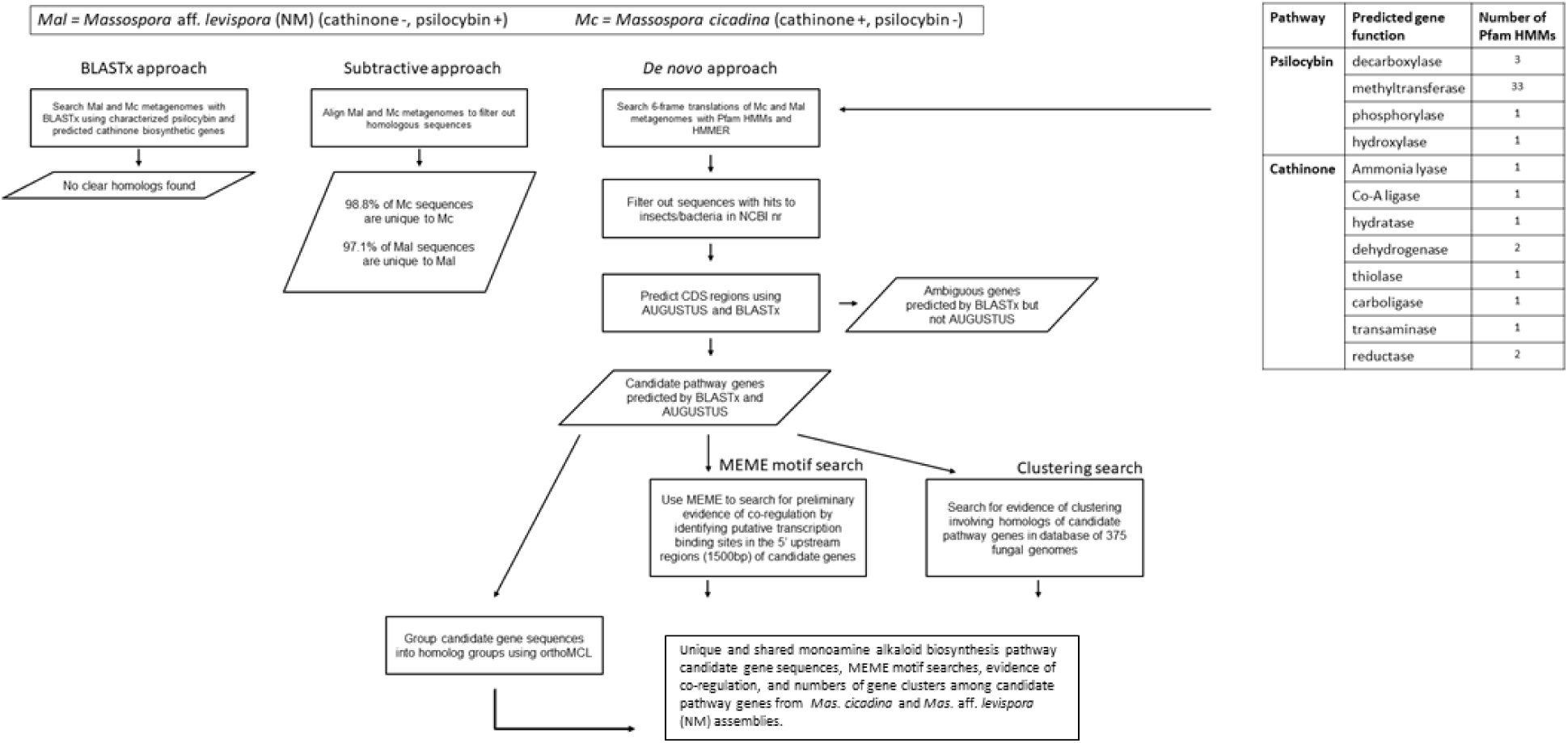
Cathinone and psilocybin biosynthesis pathway gene discovery pipeline.

**Figure 6–figure supplement 2.**
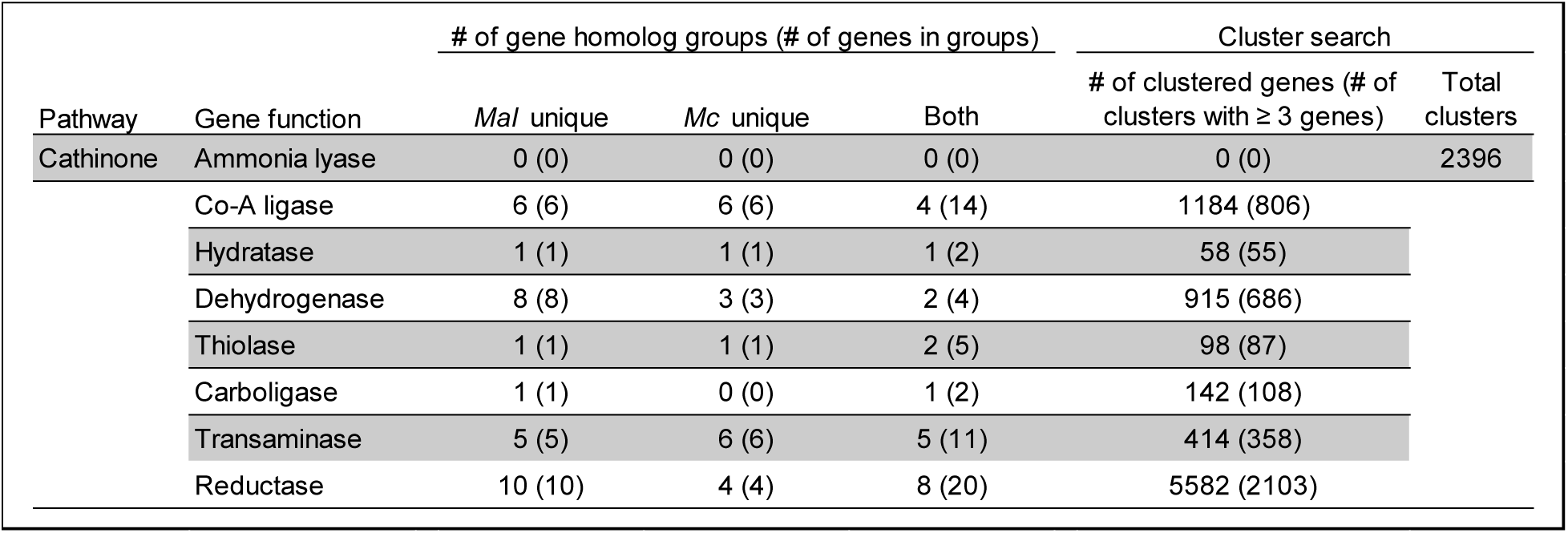
Unique and shared cathinone biosynthetic pathway candidate gene sequences, evidence of co-regulation, and number of gene clusters among containing various candidate cathinone pathway genes from the *Mas. cicadina* and *Mas*. aff. *levispora* (NM) assemblies Mas. cicadina assembly.

**Figure 6–figure supplement 3.**
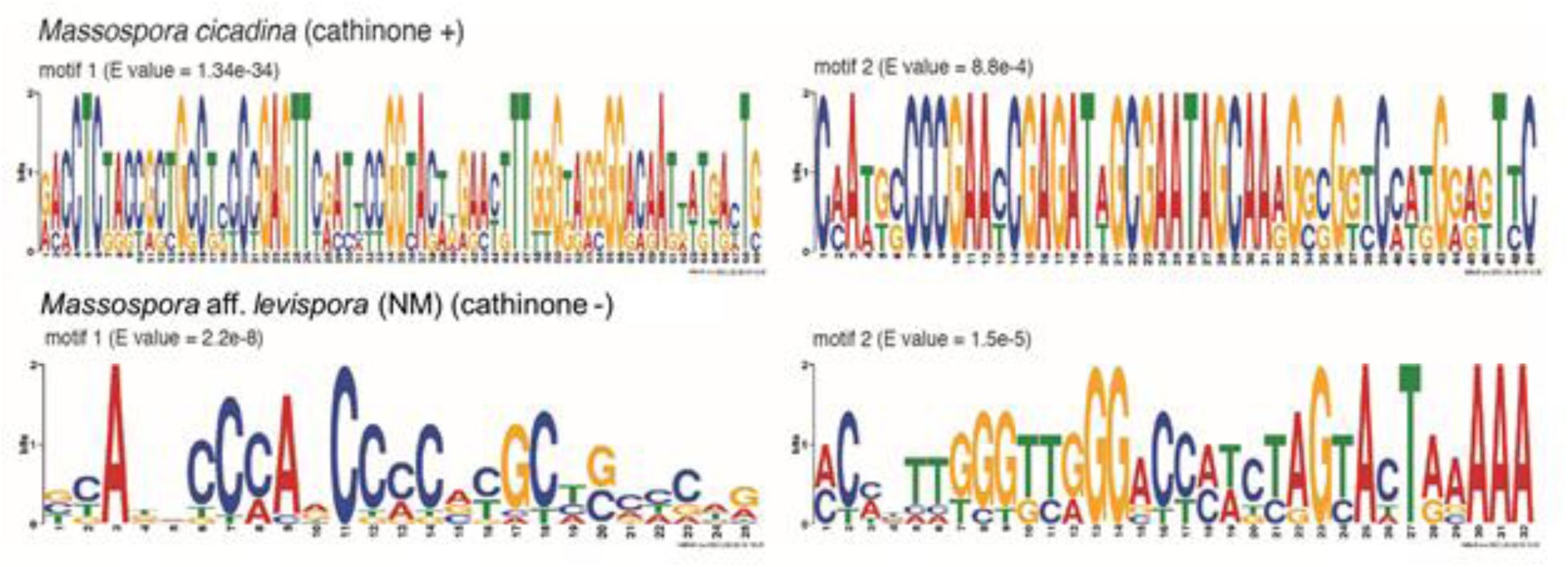
Logos of putative regulatory motifs found in the 5’ upstream regions of candidate genes in the cathinone biosynthetic pathway retrieved from the *Mas. cicadina* (cathinone +) and *Mas.* aff. *levispora* (NM) (cathinone -) assemblies. Only motifs found upstream of a predicted transaminase are shown. The median starting positions of motifs 1 and 2 from *Mas. cicadina* and motifs 1 and 2 from *Mas.* aff. *levispora* (NM) are 345bp, 405bp, 224bp and 796bp upstream of predicted translational start sites, respectively.

**Figure 6–figure supplement 4.**
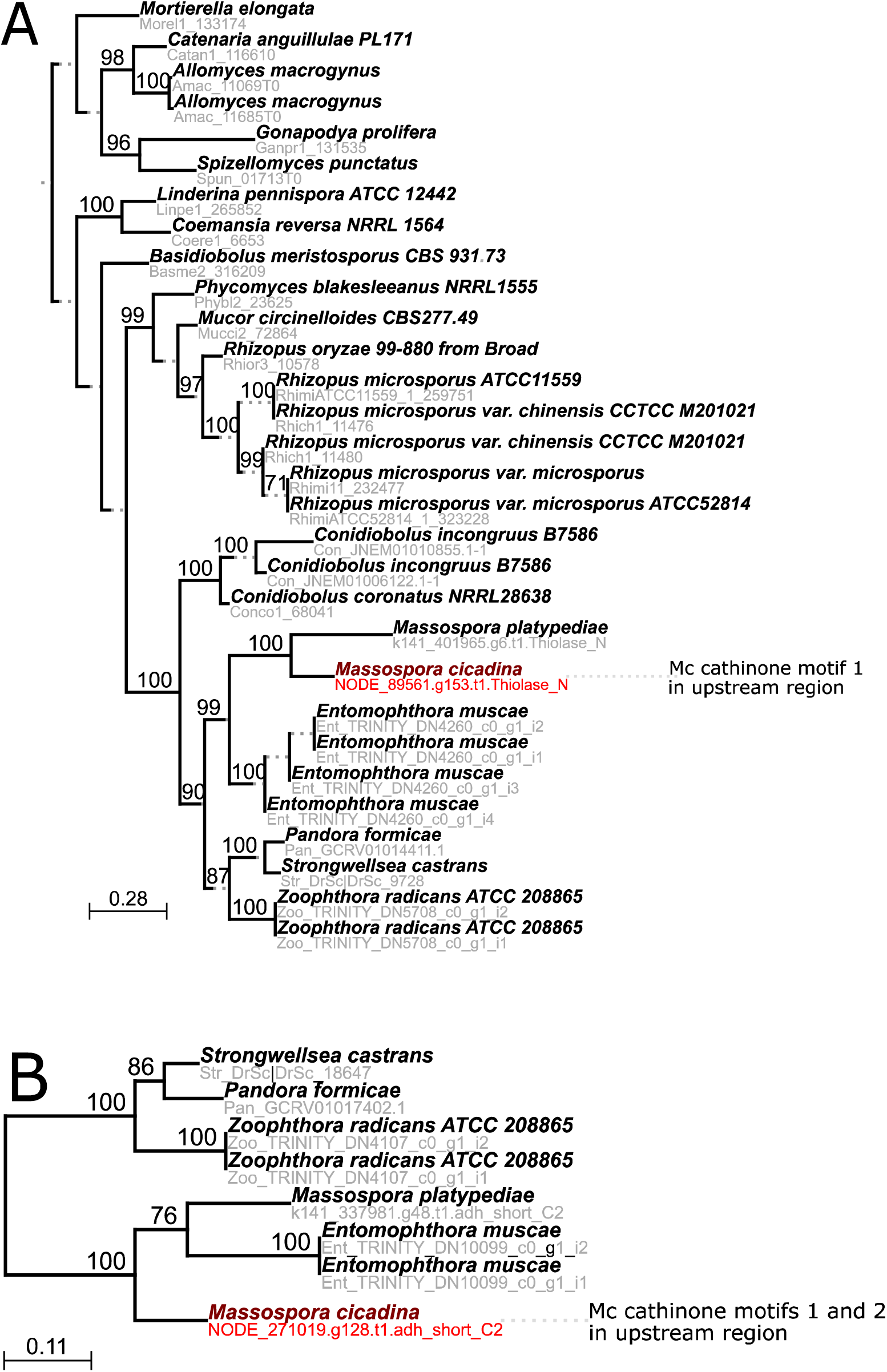
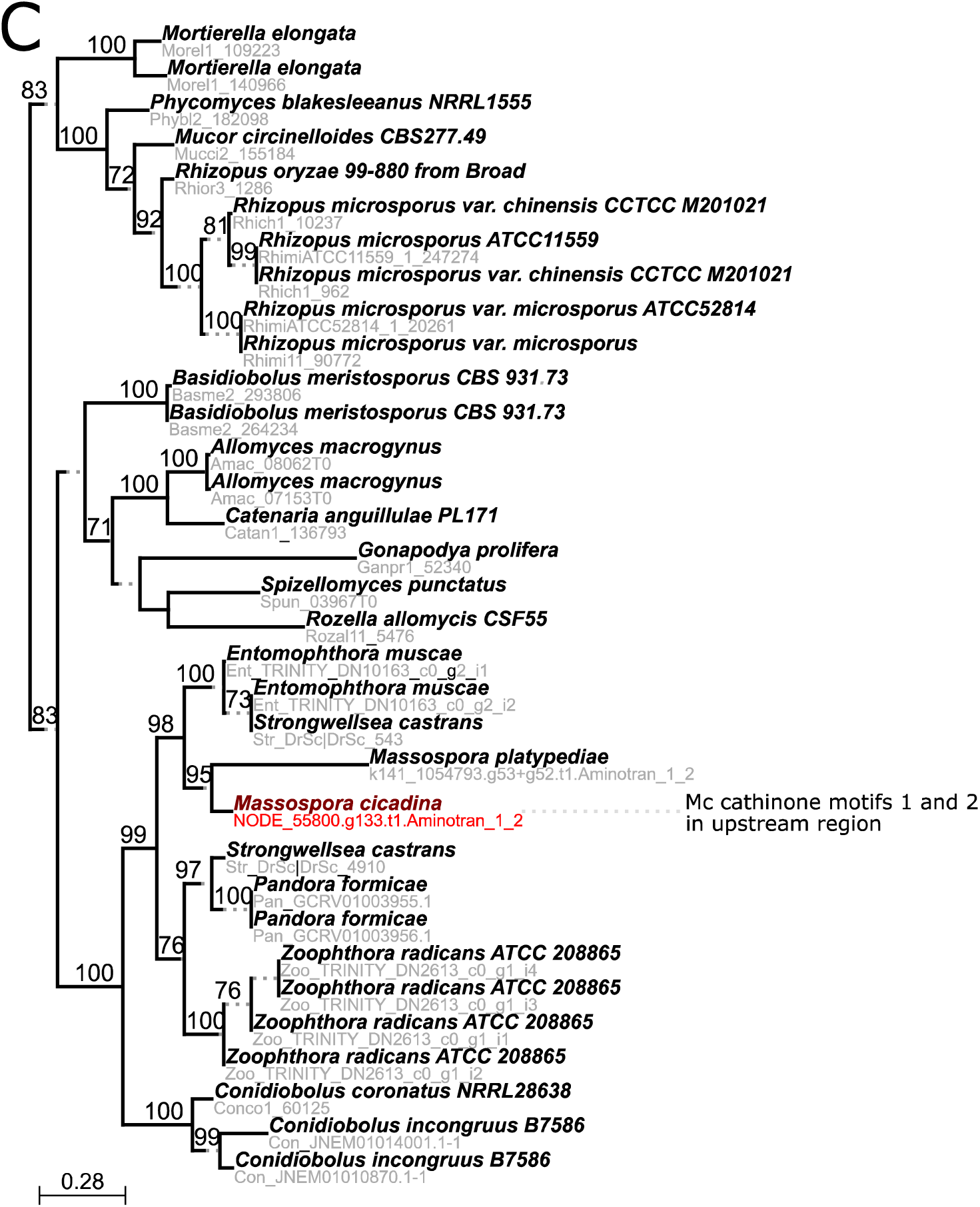
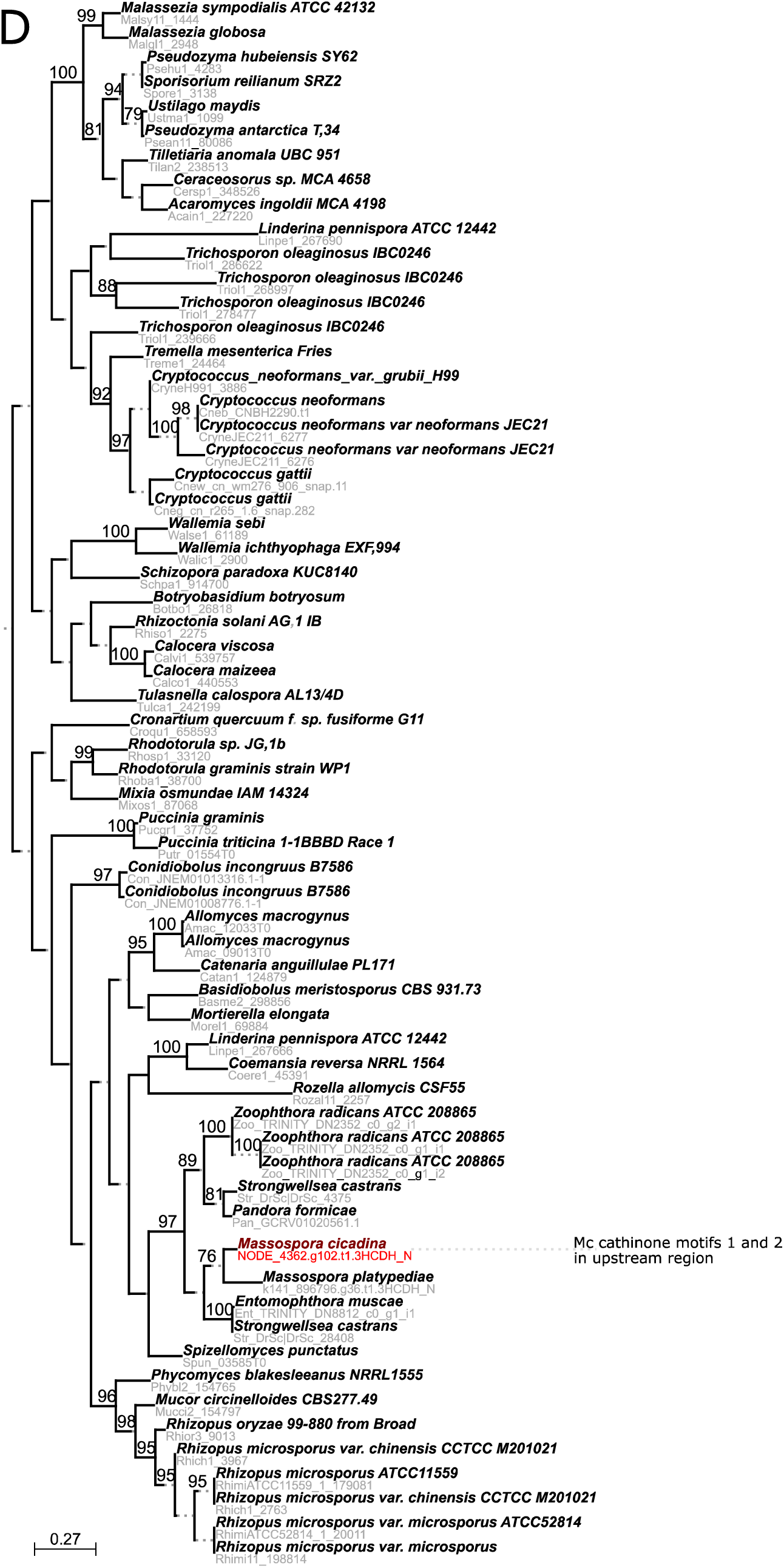
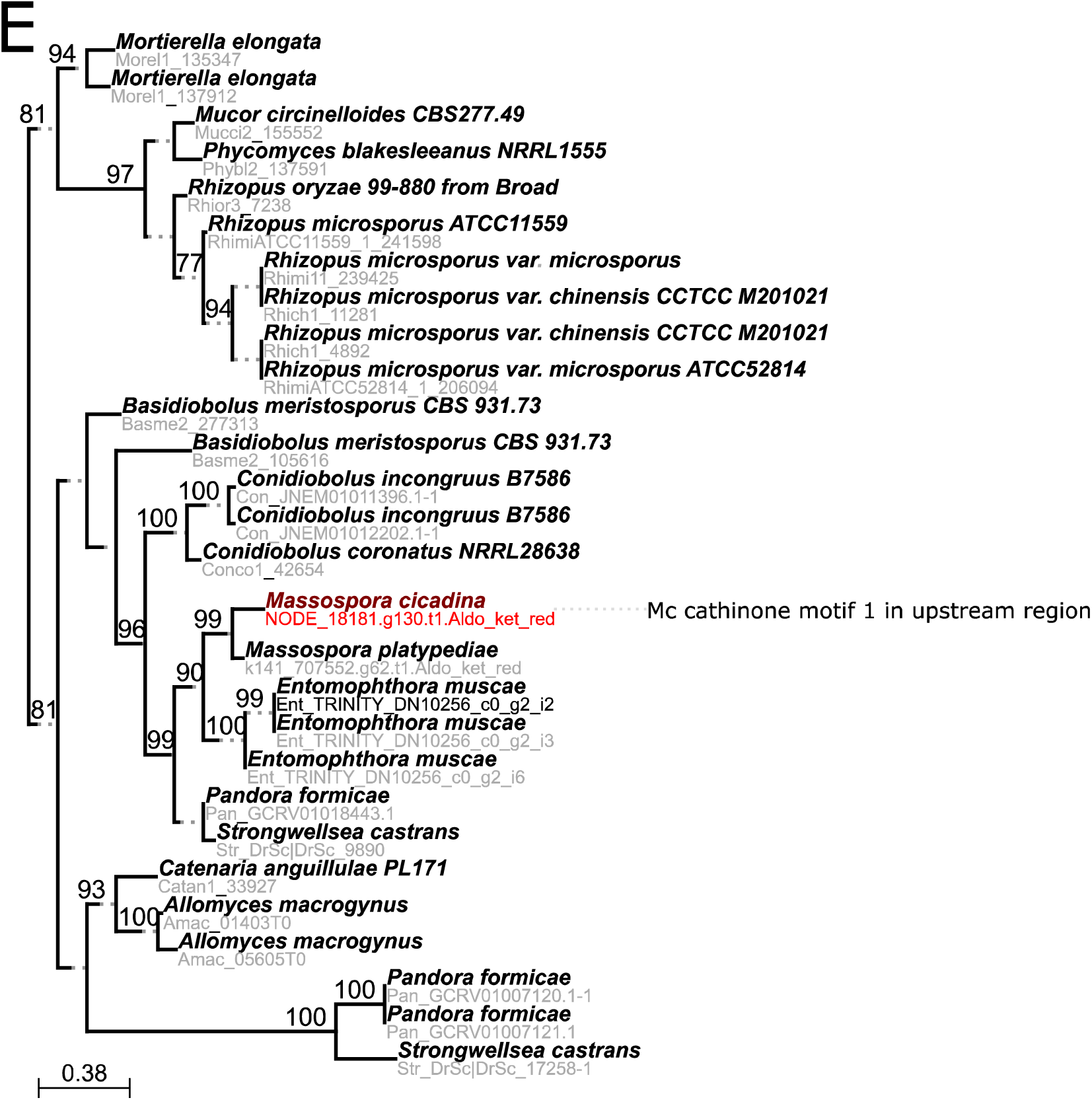
Candidate ML gene phylogenies potentially involved in cathinone biosynthesis pathway from *Mas. cicadina* and *Mas.* aff. *levispora* (NM). Phylogenies include (A) one acetyl-coenzyme A acetyltransferase (Thiolase_N), (B) one Enoyl-(Acyl carrier protein) reductase (adh_short_C2), (C) one Aminotransferase class I and II (Aminotran_1_2), (D) one 3-hydroxyacyl-CoA dehydrogenase, NAD binding domain (3HCDH_N), and (E) one Aldo-keto reductase (Aldo_ket_red).

**Figure 7–figure supplement 1.**
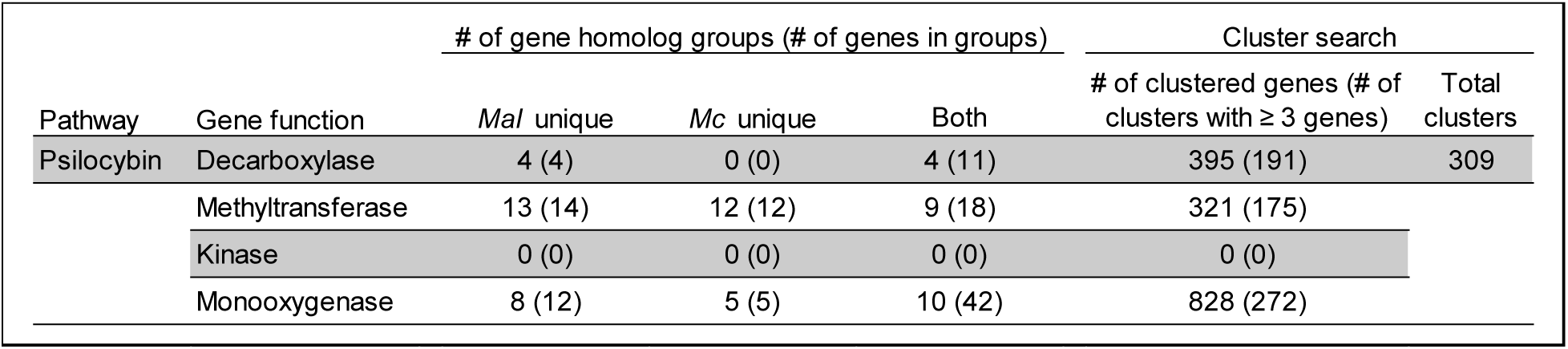
Unique and shared psilocybin biosynthetic pathway candidate gene sequences, evidence of co-regulation, and number of gene clusters among containing various candidate psilocybin pathway genes from the *Mas. cicadina* and *Mas.* aff. *levispora* (NM) assemblies

**Figure 7–figure supplement 2.**
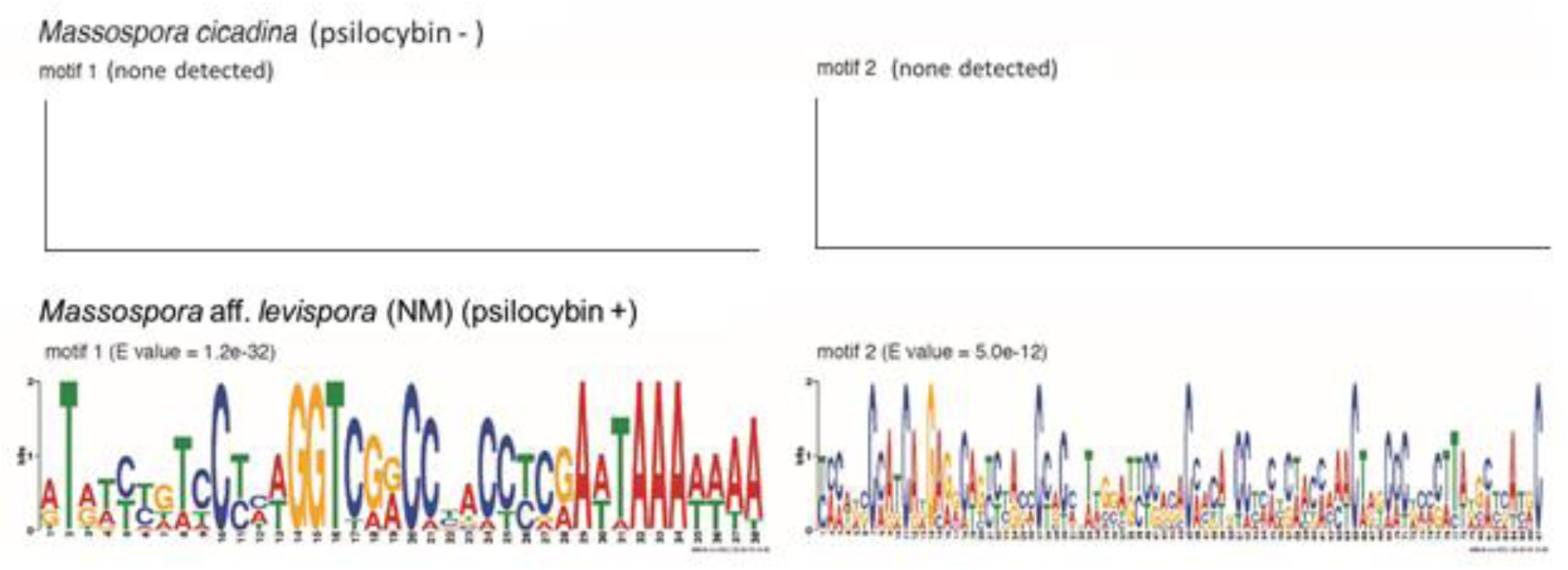
Logos of putative regulatory motifs found in the 5’ upstream regions of candidate genes in the cathinone biosynthetic pathway retrieved from the *Mas. cicadina* (psilocybin -) and *Mas.* aff. *levispora* (NM) (psilocybin +) assemblies. Only motifs found upstream of a predicted methyltransferase are shown. The median starting positions of motifs 1 and 2 from *Mas.* aff. *levispora* (NM) are 581bp and 1080bp upstream of predicted translational start sites, respectively. No putative regulatory motifs were found upstream of candidate methyltransferase genes in *Mas. cicadina*.

**Figure 7–figure supplement 3.**
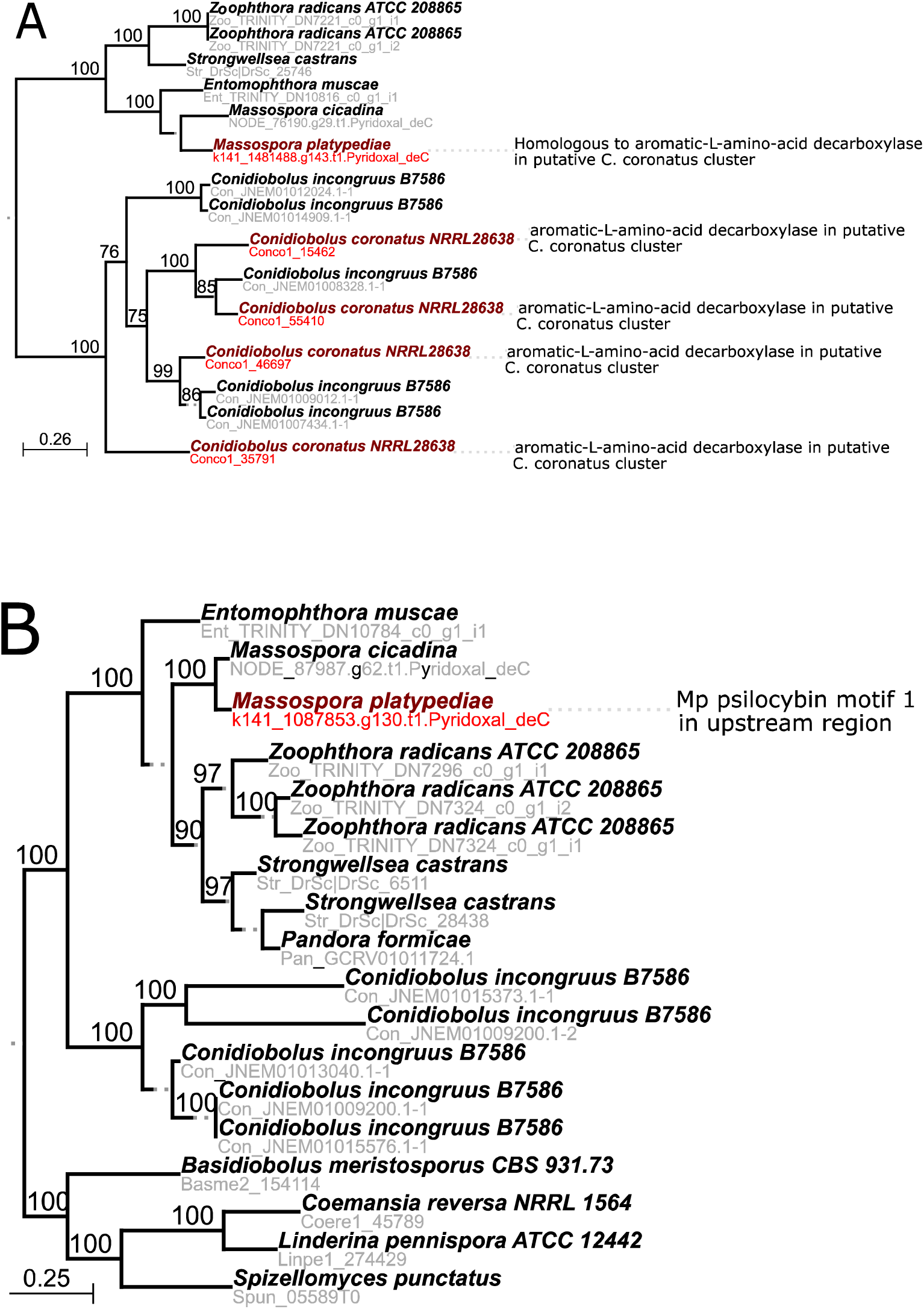
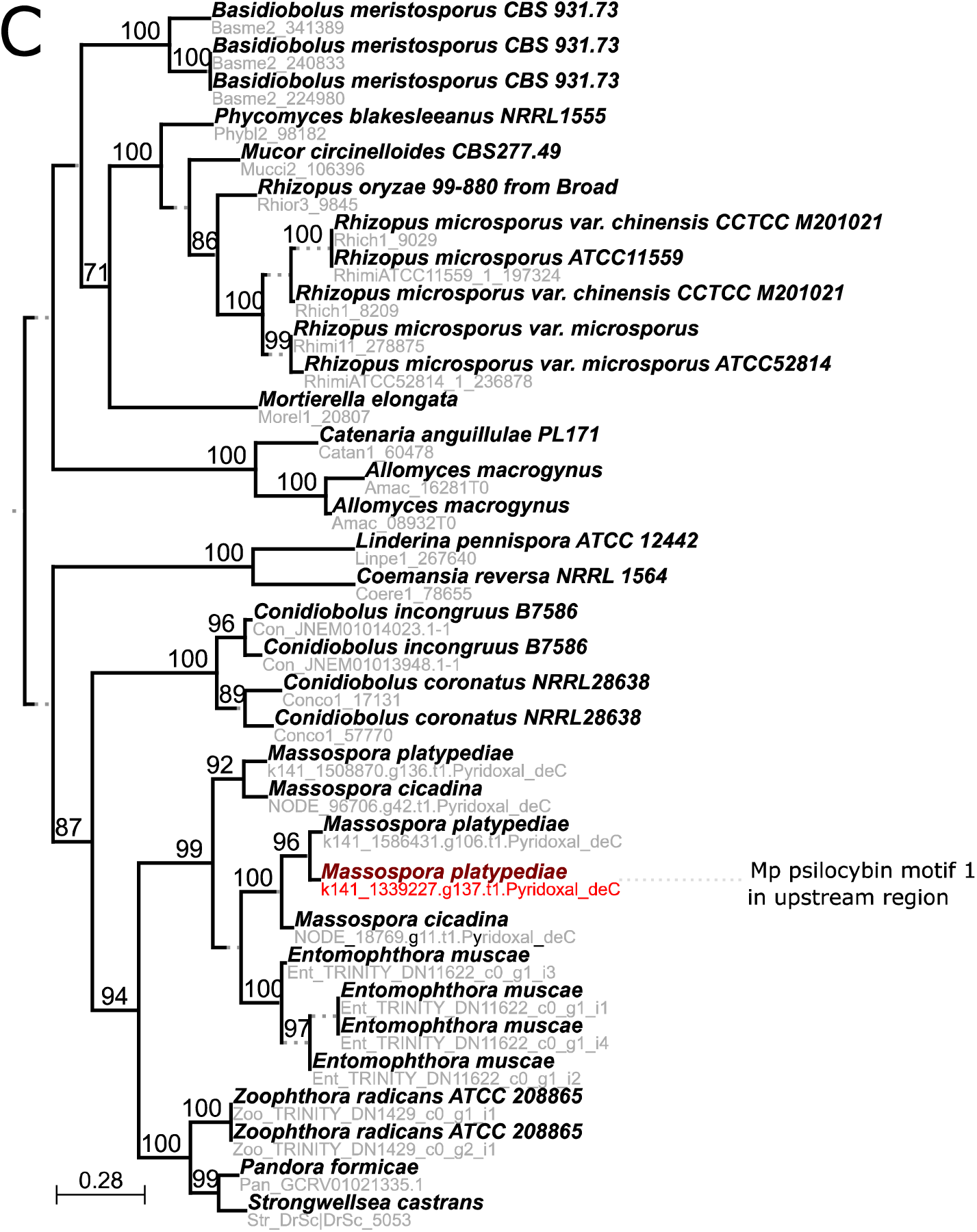
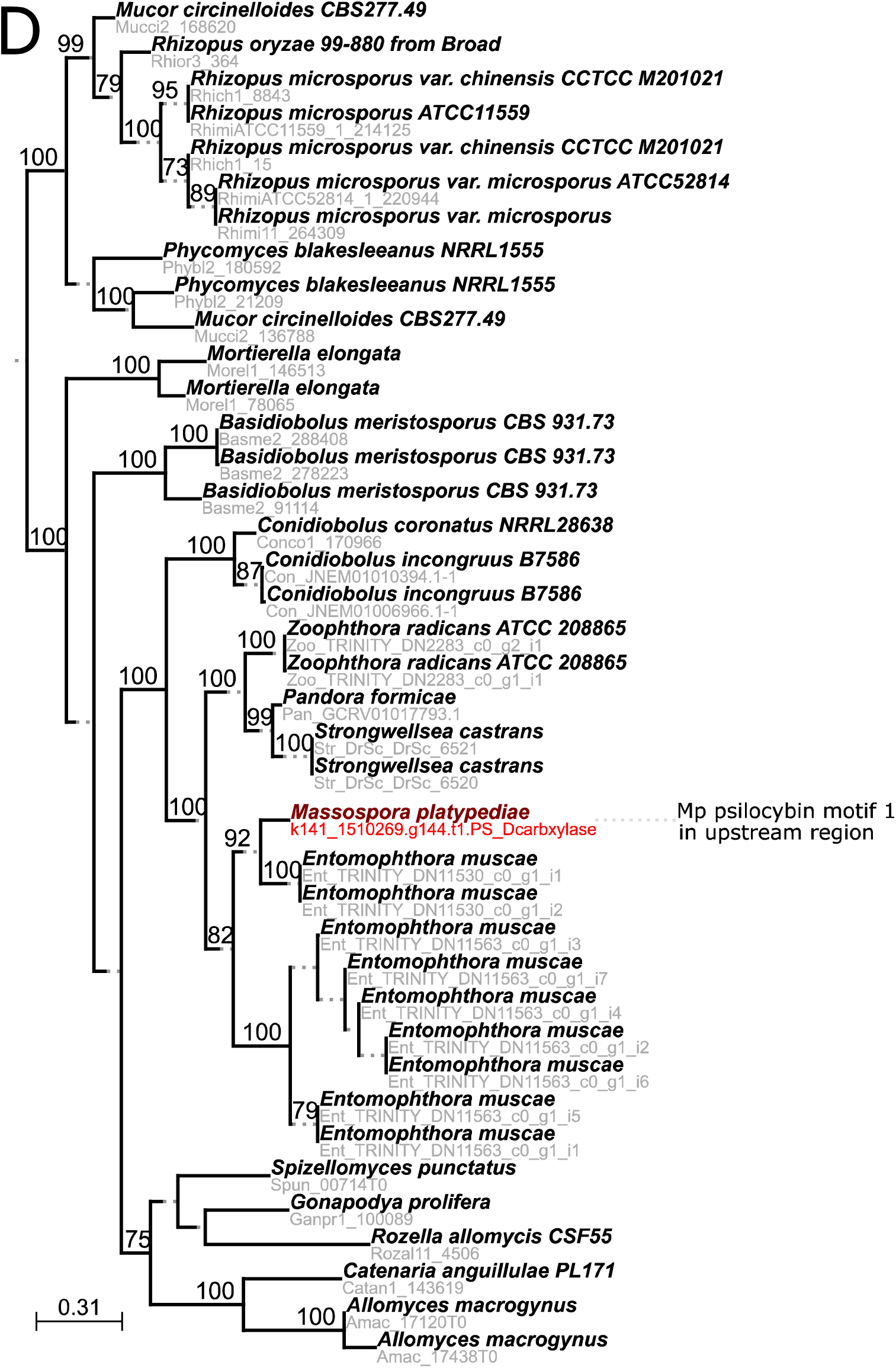
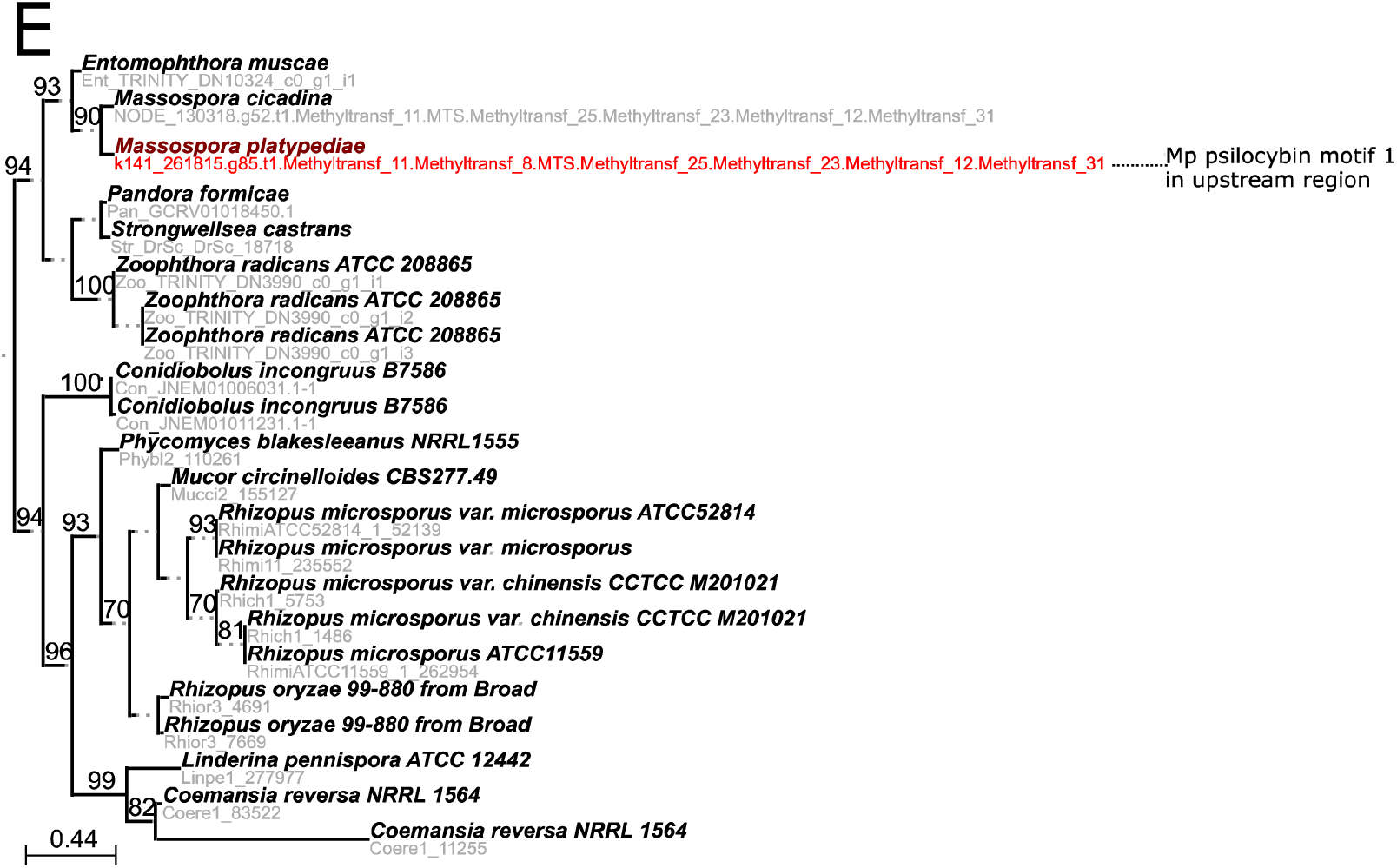
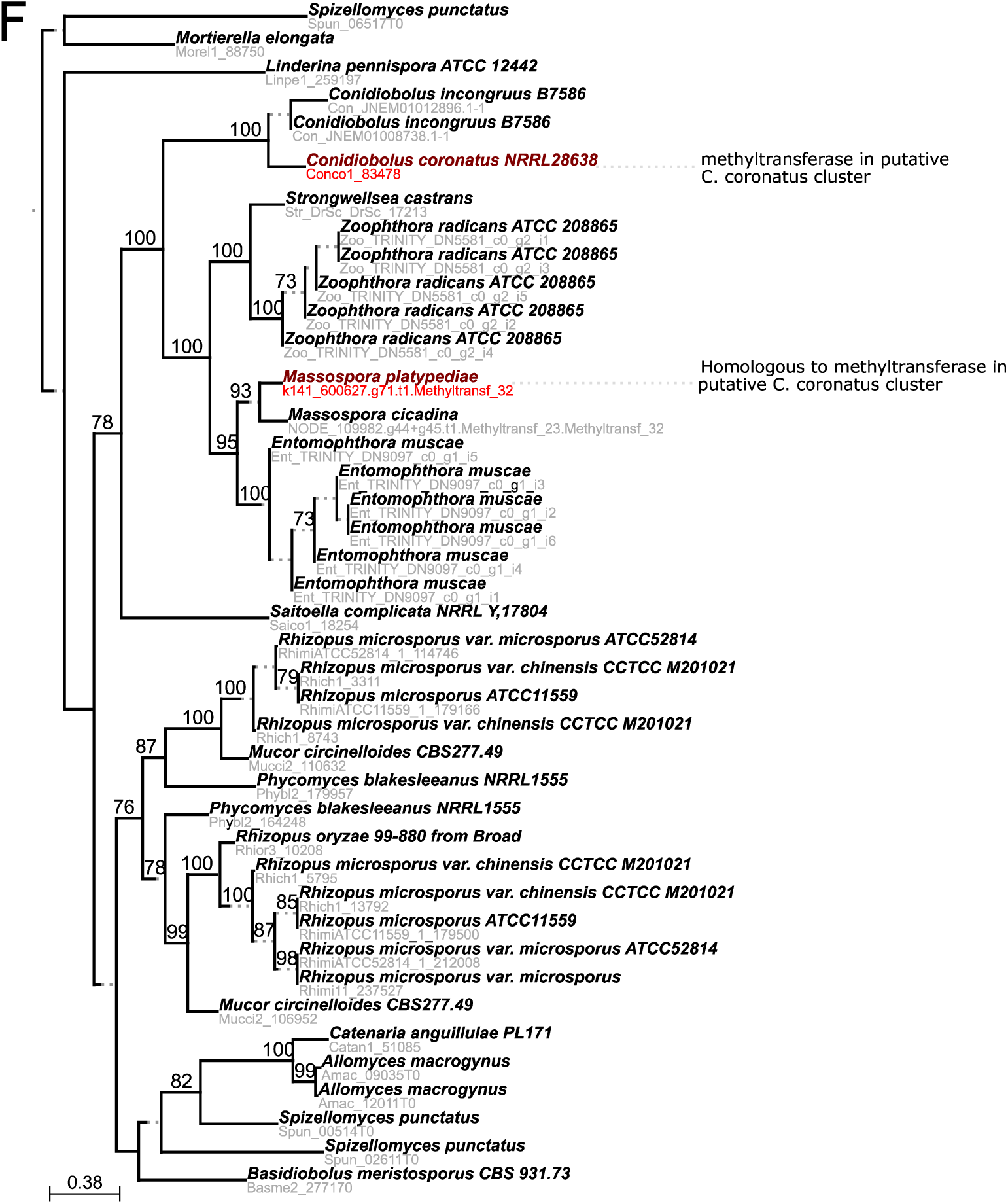
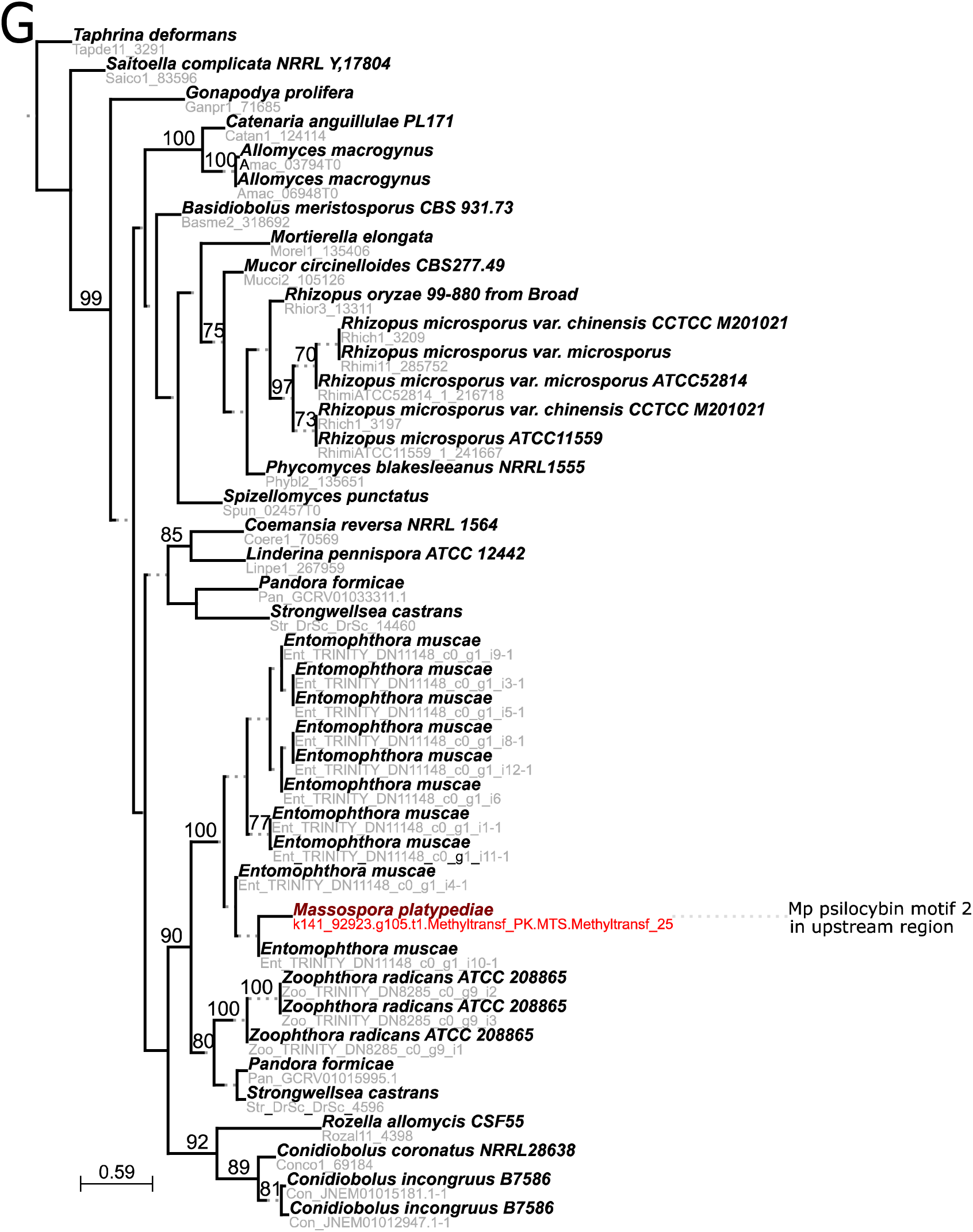
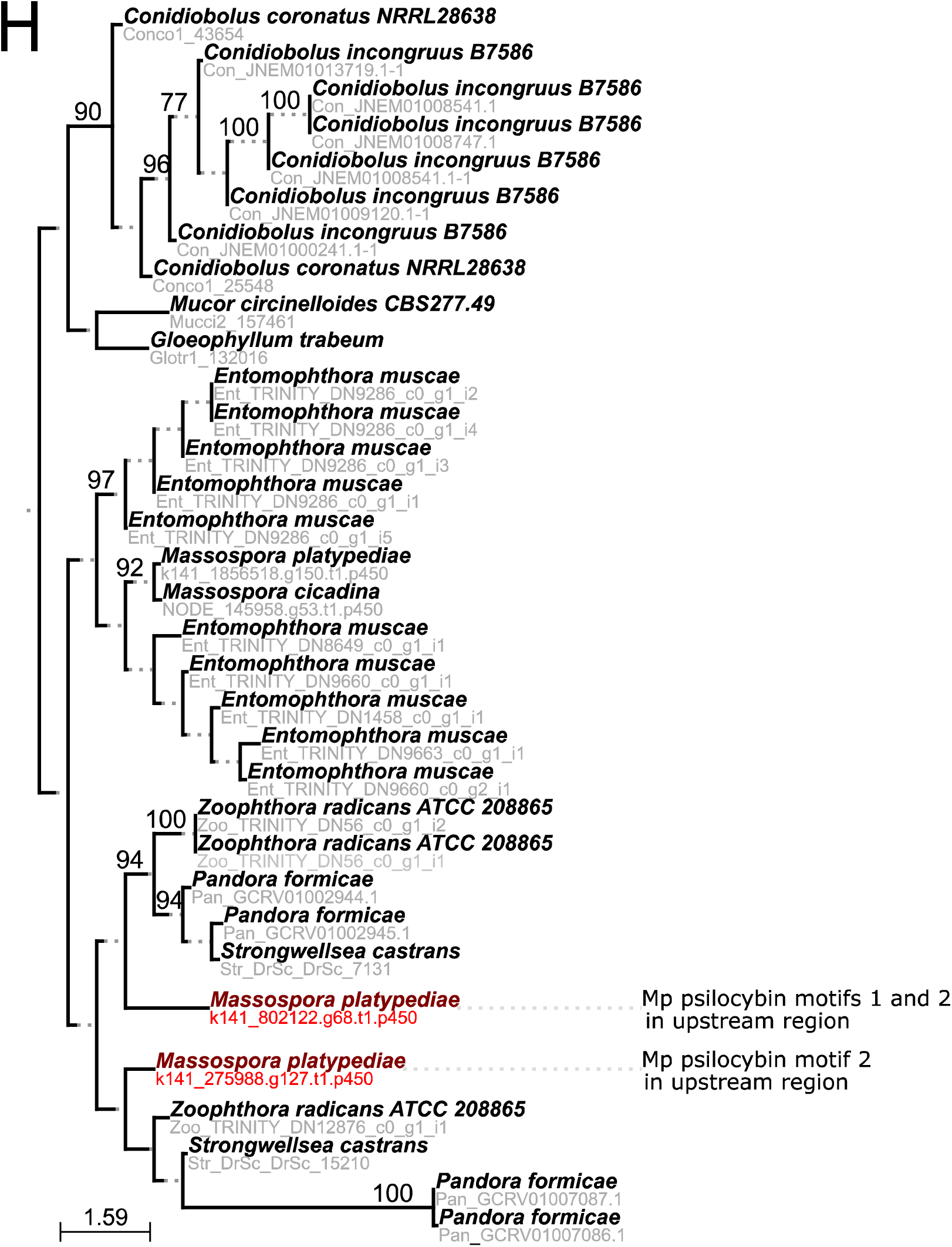
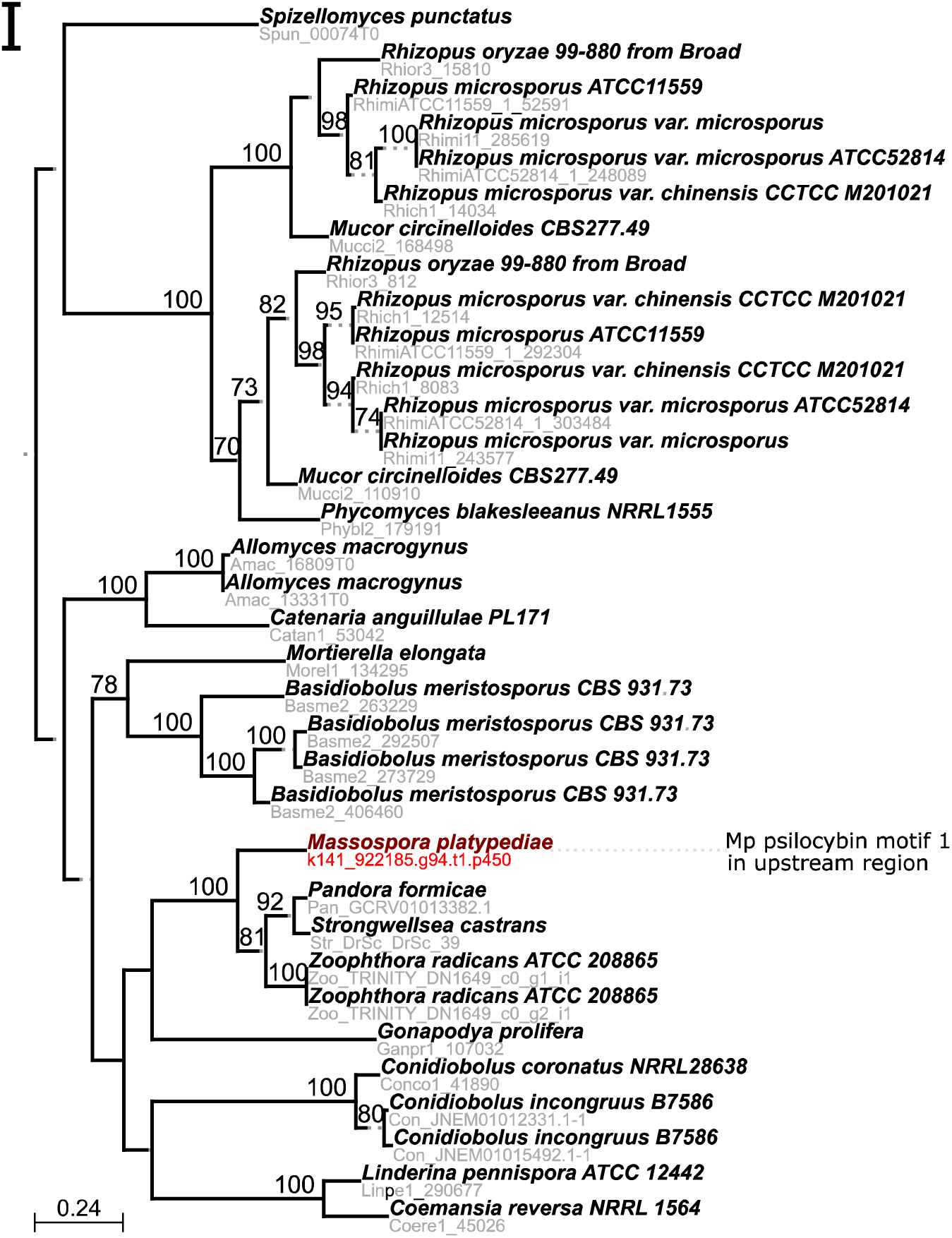
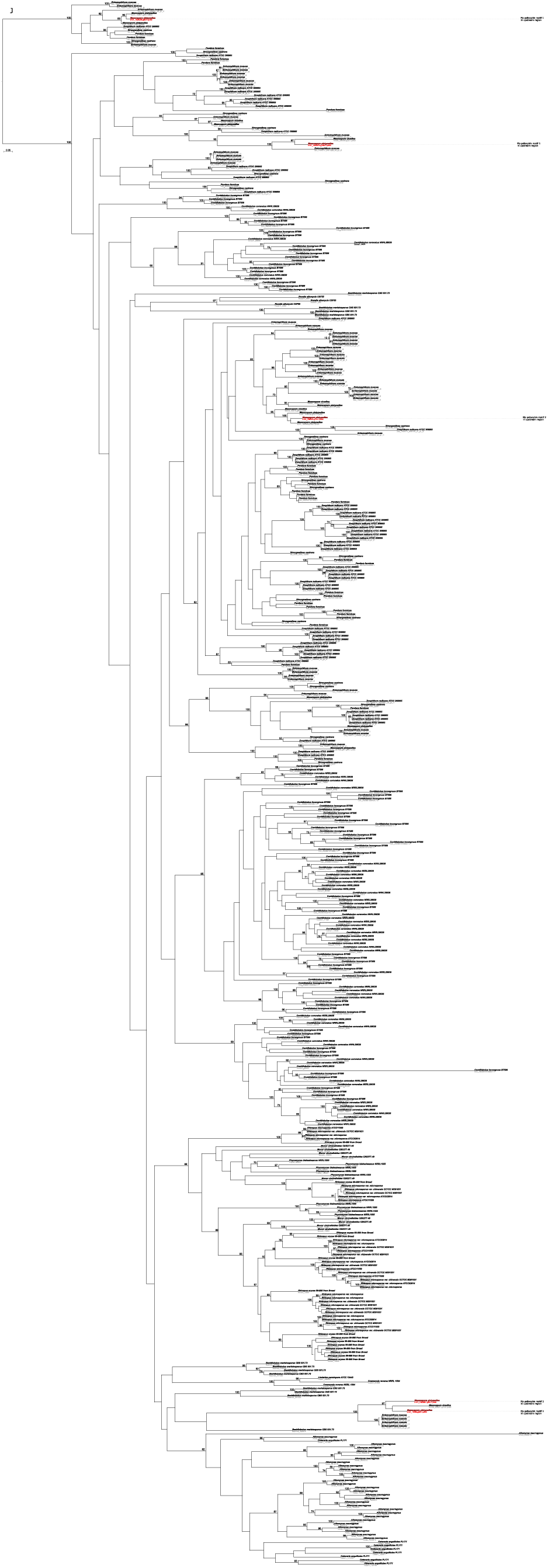
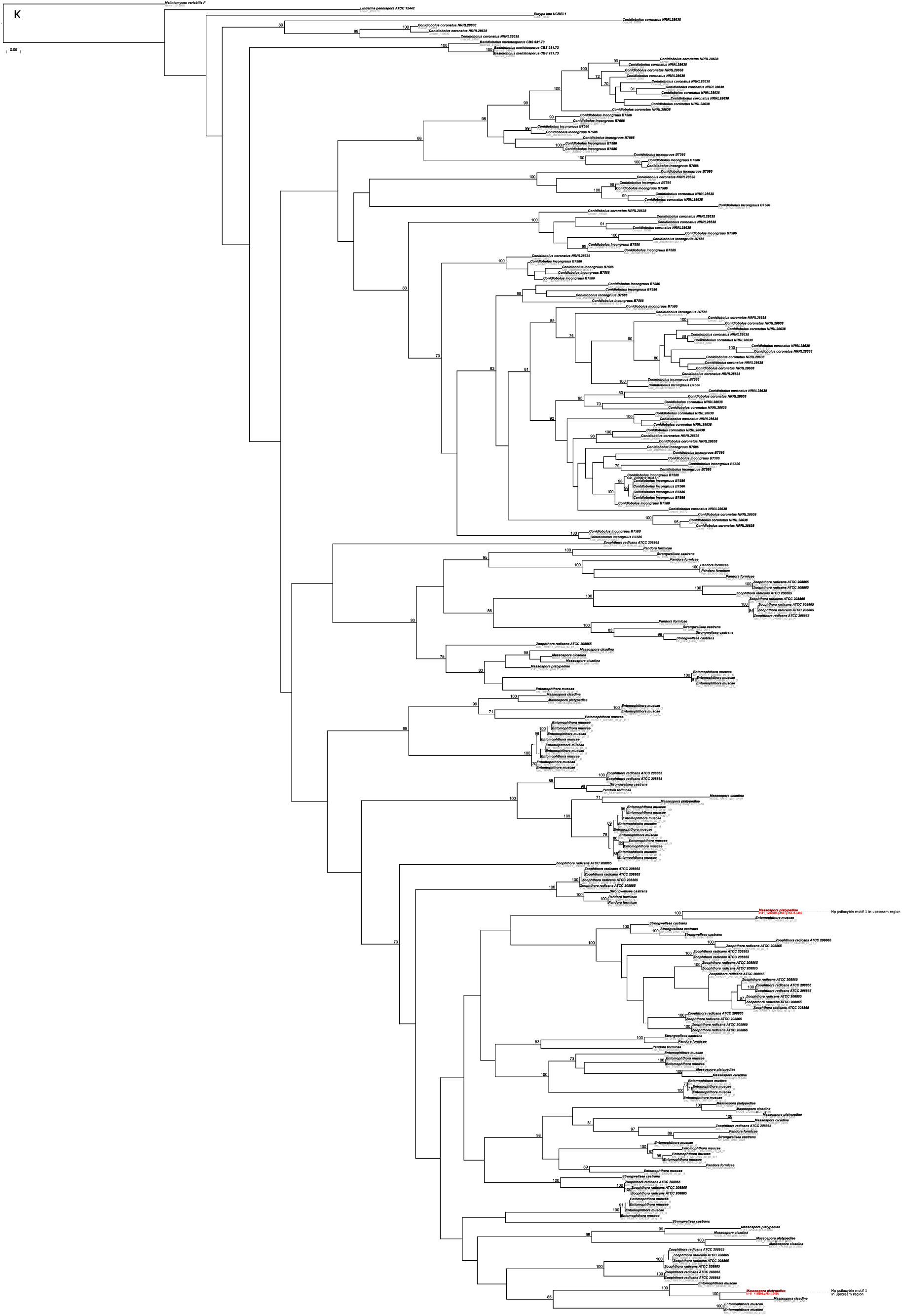
Candidate ML gene phylogenies potentially involved in psilocybin biosynthesis pathway from *Mas. cicadina* and *Mas.* aff. *levispora* (NM). Phylogenies include (A-C) three Group II pyridoxal-dependent decarboxylases (Pyridoxal_deC), (D) one Phosphatidylserine decarboxylase (PS_Dcarbxylase), (E-G) three methyltransferases (Methyltransf), and (H-K) ten Cytochrome P450 (p450).

**Figure 8–figure supplement 1.**
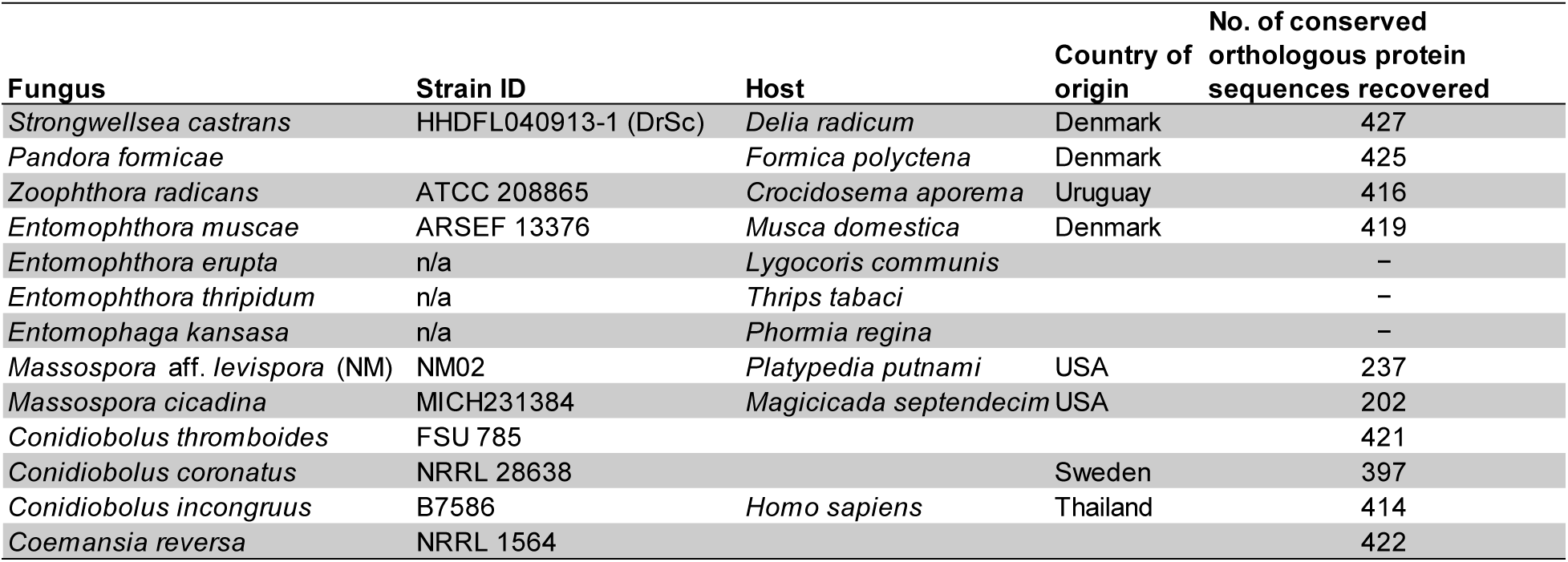
Strain histories for genome-wide phylogeny

**Figure 8–figure supplement 2.**
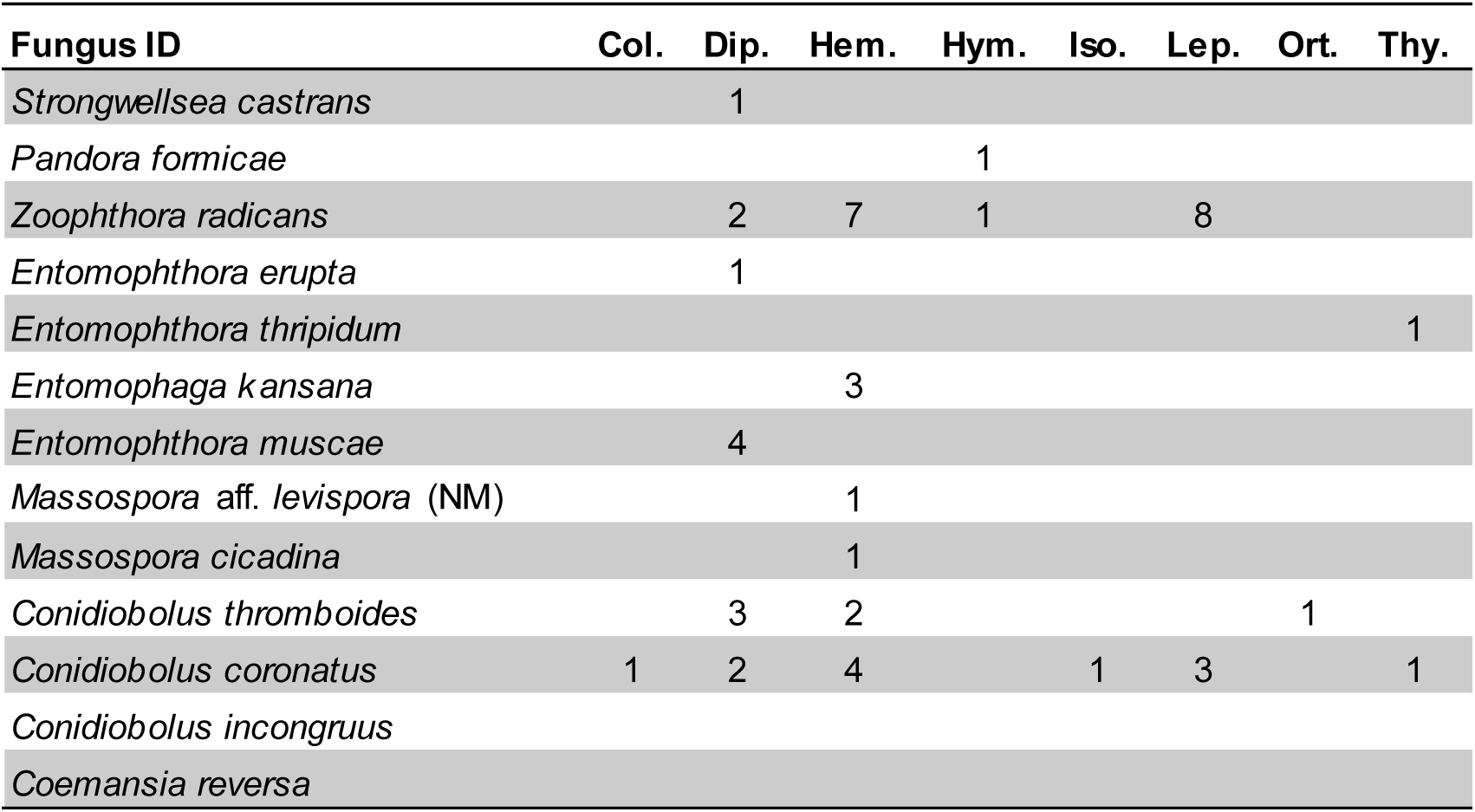
Number of susceptible insect families per order based on previously reported data from ARS Collection of Entomopathogenic Fungal Cultures (ARSEF)

**Movie 1.**
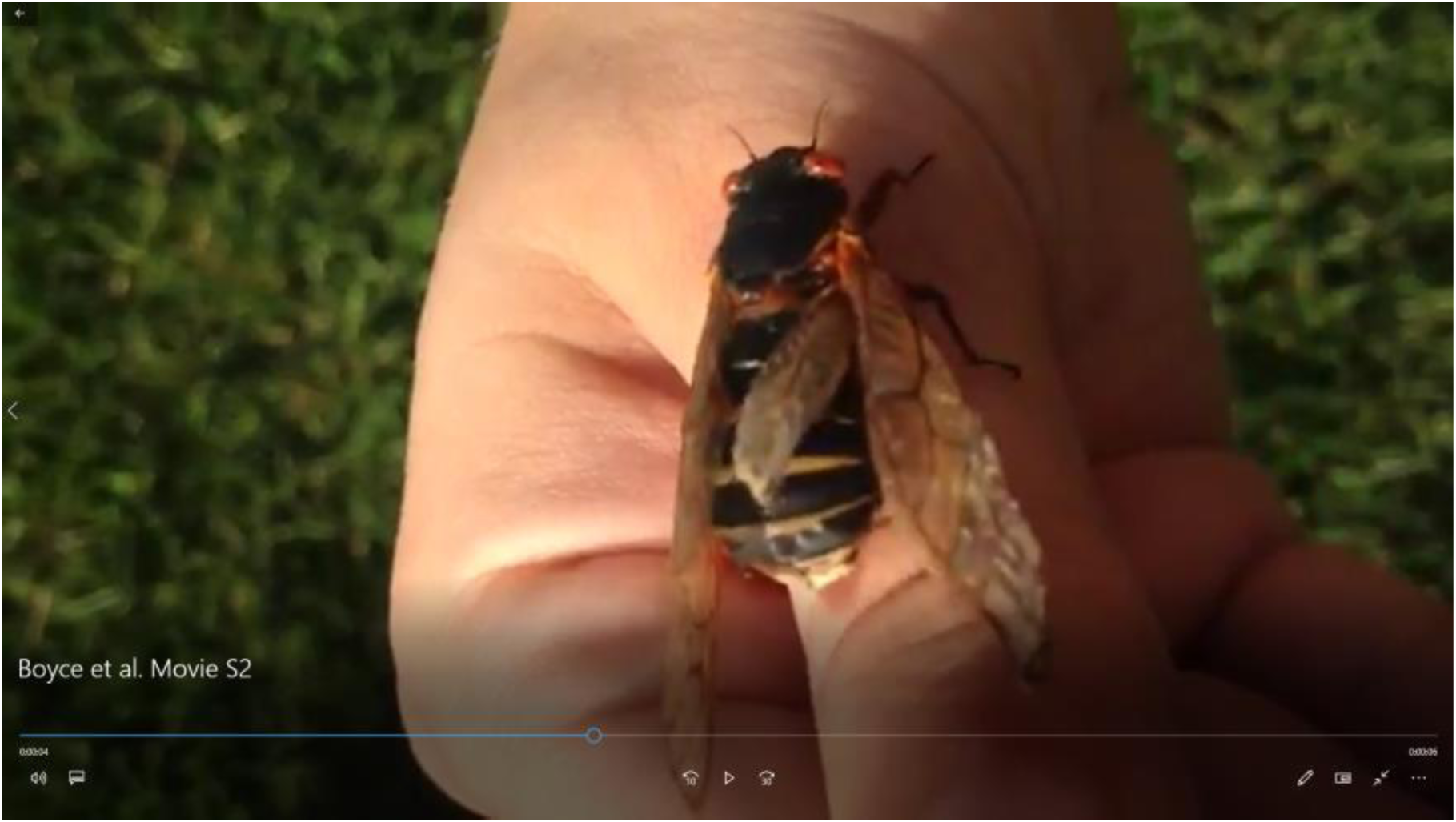
Living Brood V *Magicicada septendecim* with conidial spore infection. [This image is a placeholder for the actual movie, which will be uploaded separately. IT can be viewed here: 10.6084/m9.figshare.6855215]

Tables S1-S11 are available here: https://datadryad.org/review?doi=doi:10.5061/dryad.n9p1k0d

